# CAR-MACROPHAGES ACTIVATE ANTI TUMOR T CELLS IN THE ABSENCE OF PHAGOCYTOSIS

**DOI:** 10.64898/2026.03.11.710792

**Authors:** Lea C. Feldmann, Amaury Vaysse, Fabrice Lemaître, Béatrice Corre, Ruby Alonso, Camille Vaganay, Marion V. Guérin, Zacarias Garcia, Philippe Bousso, Capucine L. Grandjean

## Abstract

Macrophages are highly abundant within the tumor microenvironment and serve as an essential bridge between innate and adaptive immune responses. Thus, they have emerged as promising candidates for chimeric antigen receptor (CAR)-based therapeutic strategies. Previous studies demonstrated that adenovirally-transduced CAR-macrophages (CAR-M), used in clinical trials, can perform tumor cell phagocytosis, reshape the tumor microenvironment towards a proinflammatory state, and promote host T cell activation. However, how early interactions between CAR-M and tumor cells shape subsequent T cell effector functions remains poorly understood. Particularly, uncoupling the adenoviral-induced proinflammatory phenotype of CAR-M from the CAR-driven effector functions remains to be fully dissected. Here, using a CAR optimized for phagocytosis and CAR-M generated under non-polarizing conditions, we identify that CAR-M-derived cytokines and chemokines are key drivers of T cell functions while antigen cross presentation appears largely dispensable. Furthermore, dynamic imaging of CAR- M-tumor cell interactions revealed two distinct phagocytic and non-phagocytic CAR-M regardless of contact duration. Finally, we show that non-phagocytic interactions can instruct intracellular signaling and pro-inflammatory macrophage repolarization, inducing subsequent T cell activation. Altogether, these findings define an alternative mode of CAR-M action, in which CAR engagement alone, independently of target cell uptake, is sufficient to enhance T cell functions. Our study underscores the importance of non-phagocytic CAR signaling as an additional mechanism for CAR-M effector functions, opening new avenues for the design of next generation CAR for macrophage-based therapies.

**Significance statement:** Macrophages are a crucial player in the tumor microenvironment and are emerging as promising targets for cellular immunotherapy, notably through the expression of chimeric antigen receptors (CAR). CAR-macrophages (CAR-M) are currently being evaluated in clinical trials, but their precise mechanism of action remains poorly understood. The prevailing model proposes that enhanced phagocytosis by CAR- M leads to T cell activation. Here, using dynamic imaging, we uncover substantial heterogeneity in CAR- M ability to engulf tumor cells. Particularly, we show that CAR engagement, albeit lack of tumor engulfment, can produce proinflammatory mediators to activate T cells. Thus, our study identifies phagocytosis independently of CAR signaling as a complementary mode of action of CAR-M and provides a conceptual framework for the development of next generation CAR-macrophage therapies.

## INTRODUCTION

Macrophages are among the most abundant immune cell populations in solid tumors (1). Although tumor-associated macrophages often acquire immunosuppressive and pro-tumoral functions that facilitate tumor progression, they retain a remarkable degree of functional plasticity (2). In response to appropriate cues, macrophages can adopt a pro-inflammatory state, thereby promoting anti-tumor immunity and contributing to tumor regression in some contexts (3). Through their ability to fine-tune immune responses and bridge innate and adaptive immunity, macrophages act as key regulators of the tumor microenvironment (2). This functional duality has positioned macrophages at the forefront of emerging cancer immunotherapy strategies.

Initial macrophage-targeting therapeutic strategies aimed to reduce the macrophage abundance in tumors, either by selectively eliminating immunosuppressive populations (4, 5) or by limiting their recruitment, notably through blockade of the colony stimulating factor-1-receptor axis (6,7). Recently, efforts have shifted toward exploiting macrophage plasticity by reprogramming these cells toward a pro-inflammatory phenotype, by stimulating activating cascades (e.g. CD40, toll-like receptor (TLR) agonists) (8, 9). In parallel, antibody-based therapies have sought to enhance macrophage phagocytic activity by disrupting inhibitory axes, with the dual objective of triggering tumor clearance and antigen cross-presentation to T cells (10–13).

Alongside these approaches, a distinct strategy has emerged involving the engineering of macrophages to express chimeric antigen receptors (CAR-M), inspired by the clinical success of CAR-T cell therapies in hematological malignancies (14, 15). CAR-M have now undergone early-phase clinical trials, however their mechanisms of action remain incompletely understood (16). Adenovirally- generated CAR-M expressing a CAR composed of the canonical T cell signaling domain CD3ζ, have been shown to mediate tumor cell phagocytosis (17). These CAR-M acquired a strong pro-inflammatory phenotype, mainly driven by the adenoviral transduction approach, which is associated with enhanced T cell responses, (18, 19). How this pre-acquired pro-inflammatory CAR-M phenotype can be uncoupled from CAR-driven effector functions remains an unresolved question.

A current model posits that CAR-mediated engulfment of tumor cells drives antitumor immunity not only through direct tumor debulking but also by T cell priming through antigen cross-presentation (15). Under this framework, optimizing CAR-M phagocytic activity is expected to directly enhance T cell activation and therapeutic efficacy. However, whether CAR-M-mediated T cell activation strictly depends on phagocytosis or instead can be driven by alternative CAR-dependent signaling pathways, remains to be addressed.

Here, we challenge the prevailing view that the therapeutic activity of CAR-M primarily relies on tumor phagocytosis. Using real-time imaging with functional reporters of phagocytosis and CAR-induced calcium signaling, we reveal substantial functional heterogeneity among CAR-M, with many cells engaging tumor targets and activating CAR signaling without engulfing them. Despite the absence of phagocytosis, CAR engagement is sufficient to induce a pro-inflammatory transcriptional program, promote cytokine production, and enhance T cell activation. Collectively, we show that CAR engagement itself as a key instructive signal that promotes CAR-M pro-inflammatory functions, regardless of phagocytosis, thereby enhancing adaptive immune responses.

## RESULTS

### Retrovirally-transduced non-polarized CAR-M can mediate tumor control

In clinical trials, CAR-M therapies rely on adenoviral transduction of a first-generation CAR containing the CD3ζ immunoreceptor tyrosine-based activation motifs (ITAMs), a canonical T cell signaling module that can also activate Src-family kinases in macrophages (17). Mechanistic *in vivo* studies have demonstrated that these pro-inflammatory CAR-M engulf tumor cells, repolarize the tumor microenvironment and enhance anti-tumor T cell responses (18–20). In these settings, adenoviral transduction was required for polarizing CAR-M into a pro-inflammatory state, a prerequisite for therapeutic efficacy. Thus, to explore the antitumoral effect of CAR-M, we first needed to set-up a model allowing us to uncouple the consequences of adenoviral transduction from the CAR-dependent mechanism of action of the therapy. We decided to rely on the retroviral transduction of a macrophage-specific CAR-construct recognizing the model antigen CD19. This anti-CD19 CAR, composed of the intracellular FcRγ-ITAM and a PI3K recruiting domain, was shown previously to drive efficient CAR-mediated phagocytosis (17) (**Fig. 1A**). Bone marrow-derived macrophages (BMDM) were transduced with the CAR construct (CAR-M) or with empty retroviral particles (M^Empty^) as a control (**Fig. 1B, Fig. S1A-C)**. CAR-expression was assessed by surface expression of the human CD34 (hCD34) reporter encoded along the CAR construct via a self-cleaving 2A peptide or by direct CAR detection through antibody-recognition of the CAR linker (**Fig. S1D**).

**Figure 1.**
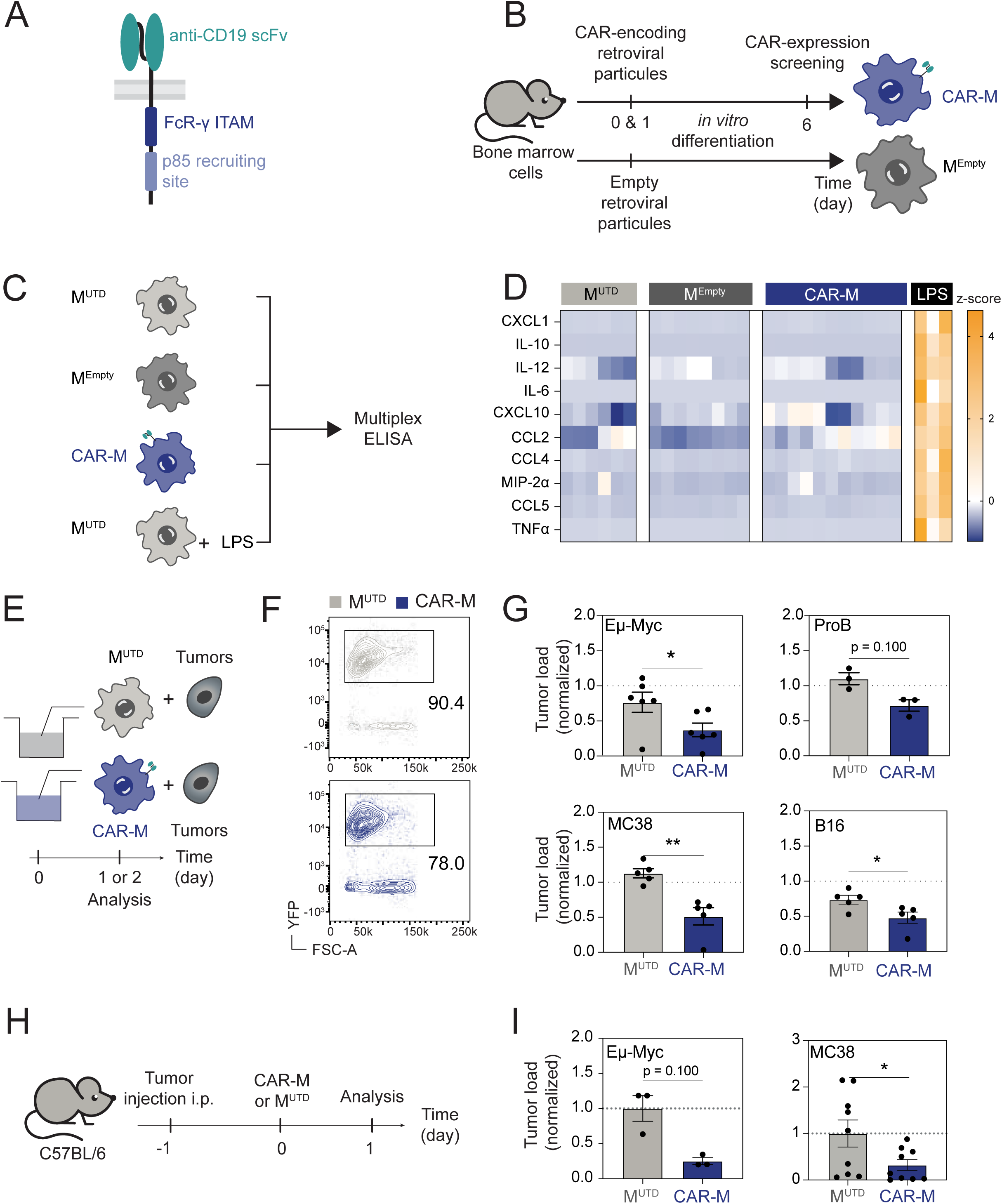
CAR-M generated by retroviral transduction are not pro-inflammatory at steady-state and reduce tumor load *in vitro* and *in vivo*. **A**. Schematic representation of the chimeric antigen receptor (CAR) expressed in macrophages. The CAR comprises an extracellular single-chain variable fragment (scFv) targeting mouse CD19, a CD8 transmembrane domain, and an intracellular domain containing the FcR-γ immunoreceptor tyrosine- based activation motif (ITAM) and the p85 subunit of PI3K-recruiting motif of murine CD19. **B.** Experimental scheme of the transduction protocol. Murine bone marrow cells were isolated, retrovirally transduced twice (day 0 and day 1), and differentiated into macrophages using L929 supernatant or m- CSF. **C–D.** Experimental design (C) and multiplex ELISA quantification of cytokines and chemokines (D) produced by untransduced macrophages (M^UTD^), sham-transduced (M^Empty^), or CAR-expressing macrophages (CAR-M), compared with LPS-stimulated untransduced macrophages. Supernatants were collected at 24 h post-stimulation and are represented as a z-score (n = 3-11 biological replicates). **E–G.** Experimental scheme (E), representative contour plots of the live cells after 24 h (Eµ-Myc) and tumor load quantification (G) following co-culture of CAR-M or control macrophages with CD19- expressing tumor cells, including two B cell tumor models (ProB and Eµ-Myc; 24 h) and solid tumors (MC38 colon carcinoma and B16 melanoma; 48 h). Tumor load was measured by flow cytometry and normalized to control tumor wells alone (n = 5–6 independent biological replicates; statistical analysis by Mann-Whitney test for not normally distributed data (Eµ-Myc, ProB and MC38) and t-test for normal distributed data (B16)). **H-I.** Experimental scheme (G) and tumor load (H) 24 h after i.p. injection of CAR-M or control macrophages (M^UTD^) into Eµ-Myc or MC38-bearing C57BL/6 mice. Tumor load was measured by flow cytometry and normalized to the control condition (n = 3–9 mice from 1-2 independent experiments; statistical analysis by Mann-Whitney test for not normally distributed data (Eµ-Myc) and t- test for normally distributed data (MC38)). Error bars represent mean ± SEM. ***P < 0.001; **P < 0.01; *P < 0.05. See Methods for additional experimental details.

As macrophage activation could be influenced by gene-transfer methods, we first determined whether retroviral transduction would tune macrophage inflammatory status. We analyzed cytokine and chemokine production over 24 h in supernatants from CAR-M and compared with untransduced macrophages (M^UTD^), M^Empty^ and macrophages pre-treated with lipopolysaccharide (LPS) as positive control (**Fig. 1C**). While LPS stimulation induced a robust inflammatory response, no substantial differences in cytokine secretion were identified between CAR-M, M^UTD^ and M^Empty^ conditions (**Fig. 1D and S2**). Thus, retroviral transduction did not intrinsically skew macrophages toward a pro-inflammatory state, and CAR-Ms generated that way do not constitutively upregulate a pro-inflammatory program at steady-state. Therefore, retroviral expression of the CAR generates largely non-polarized CAR-Ms and represents a relevant strategy to uncouple CAR-mediated effector functions from the highly polarizing adenoviral transduction approaches used in the clinic.

We next assessed the capacity of CAR-M to eliminate tumor cells both *in vitro* and *in vivo*, using hematological and epithelial/mesenchymal YFP-fluorescent tumor models. CAR-M were co-cultured with either Eµ-Myc or ProB tumor B cell lines or the CD19-engineered MC38 colon carcinoma and B16 melanoma cells all expressing comparable levels of CD19 (**Fig. 1E and S3A)**. In these conditions, we observed that CAR-M significantly eliminated all tumor cells within 48 h compared to control macrophages (**Fig. 1F-G**). This activity was strictly antigen-dependent, as demonstrated by the absence of CAR-M activity toward control CD19-negative MC38 (**Fig. S3B**). To evaluate their activity *in vivo*, we injected tumor cells intraperitoneally (using MC38, Eµ-Myc and ProB tumor cells) before treating mice with CAR-M or control macrophages. Consistent with these findings, CAR-M administration *in vivo* substantially reduced both endogenous B220^+^ B cells and tumor load, within 48h of CAR-M administration (**Fig. 1H-I and Fig. S3C-E**). Thus, non-polarized retrovirally-produced CAR- M can exert rapid antigen-specific anti-tumor activity against several tumor models *in vitro* and *in vivo*.

Altogether, these results demonstrate that regardless of a pre-acquired pro-inflammatory phenotype, CAR-M in our settings rapidly eliminate diverse tumor cell types both *in vitro* and *in vivo*, thus representing a suitable model to dissect the role of CAR signaling in driving CAR-M effector functions.

### CAR-M-derived pro-inflammatory cytokines enhance T cell activation upon CAR-M and tumor contacts

CAR-M classically used in clinical settings, have been shown to promote T cell-mediated adaptive immune responses through antigen cross-presentation, thereby contributing to sustained tumor control (18, 20). Thus, we first assessed whether in our settings CAR-M could enhance T cell activation. To this aim, CAR-M or M^UTD^ were co-cultured with ovalbumin (OVA)-expressing MC38 tumor cells, an assay ran for 4 h, similar to previous set-ups (20). This was followed by extensive washing to remove most tumor cells and the addition of CellTrace Violet (CTV)-labelled naïve OVA-specific CD8⁺ T cells (OT-I cells) for 72 h (**Fig. 2A**). We observed that in the presence of CAR-M compared to M^UTD^, OT-I T cells proliferated significantly more (**Fig. 2B**). Yet, as remaining tumor cells in the well could have impacted OT-I T cell proliferation, we decided to specifically evaluated the antigen cross-presentation abilities of CAR-M using β2-microglobulin-deficient (*β2m*^−/−^) tumor cells, preventing MHC-I surface expression (**Fig. S4A**). We observed that OT-I T cells proliferated in the presence of MHC-I-sufficient tumor cells or in presence of SIINFEKL-pulsed macrophages but failed to divide when MHC-I-deficient tumor cells were used (**Fig. 2B and S4B-C**). Thus, CAR-M enhanced T cell proliferation only in the presence of MHC-I-sufficient tumor cells (**Fig.2B**). These results suggest that initial T cell activation is driven by direct recognition of tumor antigens rather than CAR-M antigen cross-presentation. Altogether, these findings suggest that CAR-M cross-presentation is unlikely to be a major contributor of T cell proliferation.

**Figure 2.**
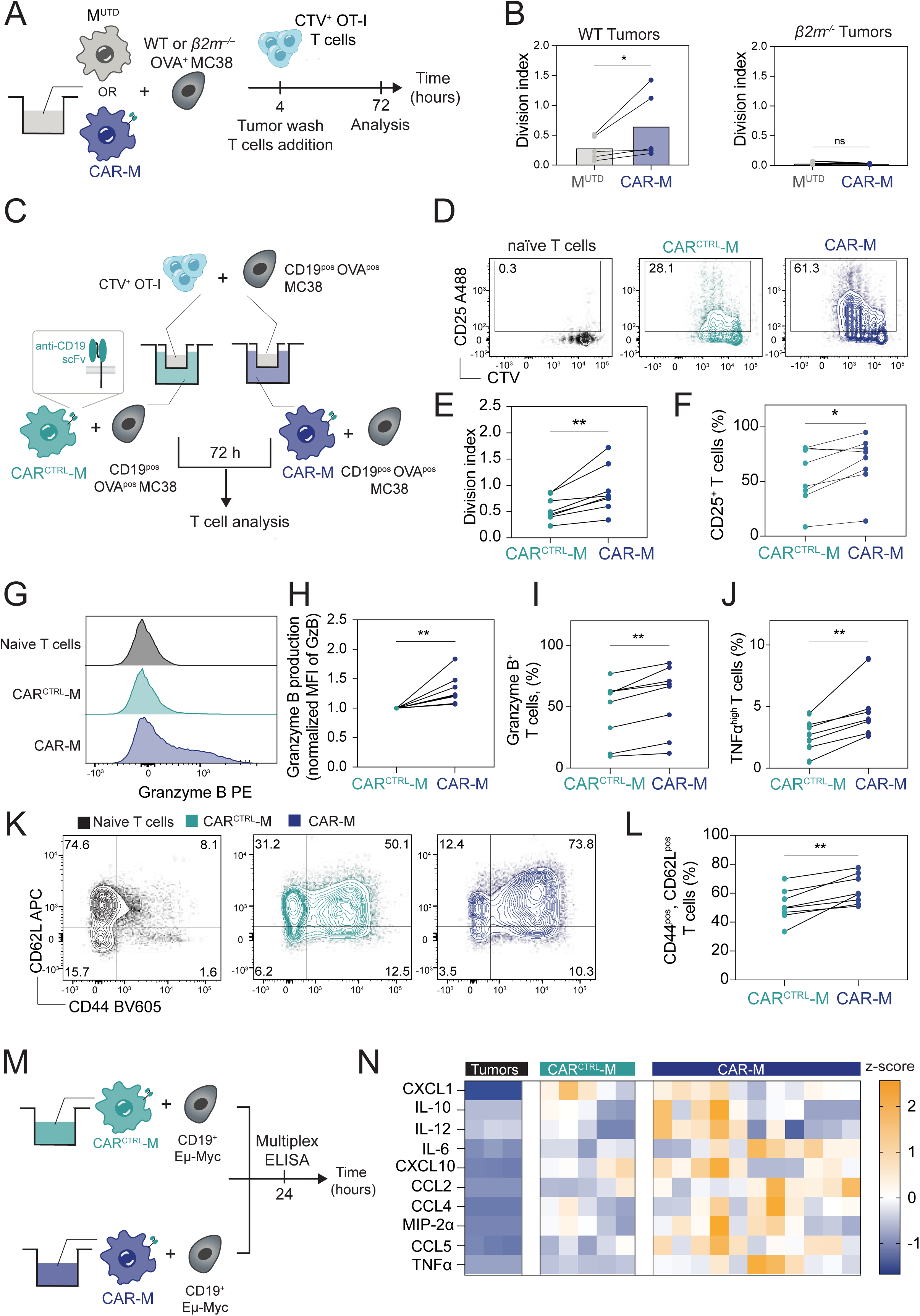
CAR-M–mediated enhancement of antigen-specific CD8^+^ T cell activation. A–B. Schematic representation of the experimental set-up (A) in which macrophages (M^UTD^ or CAR- M) were co-cultured with WT CD19⁺ OVA⁺ MC38 or *β2m*-deficient tumor cells for 4 h at an effector-to-target (E:T) ratio of 5:1. Tumor cells were subsequently washed before the addition of naïve CTV-labelled OT-I T cells at a CAR-M:T cell ratio of 5:1 and cultured in presence of IL-2 for a total of 72 h. OT-I T cell proliferation was assessed by flow cytometry and quantified using the division index (B) (n = 3–5 independent biological replicates from 5 independent experiments. Statistical analysis by ratio-paired t-test for normally distributed data (WT) or Wilcoxon test for not normally distributed data (*β2m^-/-^ )*). **C-F**. Schematic of the transwell experimental set-up (C) in which macrophages (CAR^CTRL^-M, or CAR- M) and CD19⁺ OVA^+^ MC38 tumor cells were plated in the lower chamber, while naïve OT-I T cells and OVA⁺ CD19^+^ MC38 tumor cells were placed in the upper chamber of the transwell assay. T cells were analysed by flow cytometry after 72 h in presence of IL-2, as shown in representative flow cytometry profiles (D). Quantification of the T cell division index (E) and the proportion of activated CD25⁺ T cells (F) are shown. (n = 8 independent experiments, statistical analysis by paired t-test). **G.–I.** Granzyme B expression by OT-I. G. Representative histograms of the expression of Granzyme B. Quantification of the expression levels of Granzyme B following the transwell assay, shown as mean fluorescence intensity (MFI) normalized to CAR^CTRL^-M (H) and as percentage of positive of OT-I (I) cells (n = 8 independent experiments, statistical by Mann-Whitney test (H) or paired t-test (I)). **J.** Proportion of TNFα^high^ OT-I T cells following the transwell assay (n = 7 independent experiments; statistical analysis by paired t-test). **K–L**. Representative flow cytometry profiles (K) and quantification (L) of CD44 and CD62L expression on activated OT-I T cells following the transwell assay with CAR-M or CAR^CTRL^-M (n = 8 independent experiment, statistical analysis by paired t-test). **M-N.** Experimental scheme (M) and multiplex ELISA quantification of cytokines and chemokines (N) produced when CAR^CTRL^ transduced macrophages (CAR^CTRL^-M) or CAR-expressing macrophages (CAR-M) were co-cultured with tumor cells (Eµ-Myc). Data are shown as a z-score and are from n = 5-11 independent experiments. Supernatants were collected at 24 h post-stimulation. Error bars represent mean ± SEM. ***P < 0.001; **P < 0.01; *P < 0.05. See Methods for additional experimental details.

Whether CAR-M enhanced T cells responses in a contact-independent manner for instance through soluble factors remained to be addressed. Thus, to address this question, we performed transwell assays in which CAR-M were co-cultured with tumor cells (MC38 or Eµ-Myc) and physically separated from naïve OT-I T cells cultured with CD19^+^OVA^+^ MC38 tumor cells to maintain antigen recognition (**Fig. 2C and S5A**). Under these conditions, T cells exposed to CAR-M-derived soluble factors exhibited increased proliferation, numbers and activation as shown by elevated CD25 expression only in the presence of a full signaling CAR construct and in the presence of the CD19 antigen. Indeed, T cells did not proliferate when exposed to macrophages expressing a non-signaling CAR, lacking intracellular domains (CAR^CTRL^-M) or for those exposed to CAR-M cultured with tumor cells lacking CD19 expression (**Fig. 2D-F** and **S5A-F**). These findings indicate that CAR engagement and downstream signaling in tumor-exposed CAR-M promote the release of soluble mediators that enhance T cell activation. Further functional analyses revealed that CAR-M-conditioned factors also promoted T cell cytotoxic differentiation, as indicated by increased frequencies of granzyme B^+^ T cells and elevated granzyme B expression levels (**Fig. 2G-I and S5G**). Moreover, we observed a higher proportion of tumor necrosis factor alpha ^high^ (TNF-α^high^) T cells (**Fig. 2J and S5H**). Notably, T cells exhibited a trend toward a central memory-like phenotype (CD62L^+^ CD44^+^), which has been associated with improved anti-tumor responses (21) (**Fig. 2K-L**). Together, these results indicate that CAR-M activated upon tumor antigen recognition release soluble factors that enhance T cell activation, drive effector differentiation, and bias T cells toward a central memory-like state.

To define the soluble mediators induced by CAR-M in the presence of tumor cells, we performed a multiplex cytokine and chemokine assay on supernatants from CAR-M co-cultured with Eµ-Myc or CD19^+^MC38 tumor cells and compared them to CAR^CTRL^-M (**Fig. 2M and S6A**). These co-cultures exhibited high production of pro-inflammatory cytokines and chemokines, including TNFα, CCL5, CXCL1 and IL-12, which appeared stronger with Eµ-Myc tumor cells in the first 24 h (**Fig. 2N and S6B-D**). To further evaluate whether these results translated *in vivo*, we analyzed peritoneal lavages from Eµ-Myc-bearing mice injected intraperitoneally and treated with CAR-M or M^UTD^ (**Fig.S7A- C).** Consistent with the *in vitro* data, CAR-M treatment led to increased levels of several cytokines such as TNFα or type-I interferons (IFN), as well as CCL2 and CCL4 chemokines which are implicated in T cell recruitment (22) in the first 24 h. Thus, these findings demonstrate that CAR-M rapidly start secreting cytokines and chemokines upon contact with tumor cells both *in vitro* and *in vivo.* Collectively, these findings demonstrate that CAR-M in contact to tumor cells, can enhance distal T cell activation not through antigen cross-presentation, but rather by creating a pro-inflammatory cytokine milieu, responsible in turn for promoting T cell activation and proliferation.

### CAR-M–tumor interactions result in phagocytic and non-phagocytic outcomes

As the presence of tumor cells is required in our model for CAR-M to adopt a pro-inflammatory phenotype, we next sought to dissect the outcome (e.g. phagocytosis or apoptosis) of early CAR-M tumor cell interactions. To monitor phagocytosis and apoptosis, tumor cells were engineered to express a dual-fluorescent Förster resonance energy transfer (FRET) reporter, enabling simultaneous quantification of both processes by flow cytometry and dynamic imaging (23) (**Fig. 3A**). Briefly, live cells appear FRET positive due to energy transfer between CFP and YFP proteins held in proximity by the short caspase-3 cleavable DEVD peptide. During apoptosis, caspase-3-dependent cleavage of DEVD abrogates CFP-YFP interactions resulting in FRET loss. During phagocytosis, the acidic phagolysosomal compartment drives the progressive loss of YFP fluorescence (YFP^low^, blue), while CFP signal is retained, also leading to a FRET loss only once the cell is engulfed. Thus, this probe allows us to quantify phagocytic and cytotoxic events in co-cultures of CAR-M and tumor cells (23). To investigate the outcomes of CAR-M and tumor interactions, we first quantified phagocytosing macrophages in co-cultures of CAR-M with different CD19-expressing tumor cells (**Fig. 3A**). During flow cytometry, phagocytosing macrophages were defined as F4/80⁺ CAR-bearing macrophages exhibiting low YFP expression. Unexpectedly, although the proportion of phagocytosing macrophages varied across tumor types, phagocytic CAR-M accounted for no more than 35% of the CAR-M population over 1 h and occurred only in the presence of CD19-expressing tumor cells (**Fig. 3B and S8A**). To further clarify CAR-M–tumor interaction dynamics, we next performed real-time imaging *in vitro* and visualized FRET^+^ (YFP^+^ and CFP^+^; magenta), and FRET^−^ (YFP^+^; blue) tumor cells (**Fig. 3C, S9 and Movie S1**). While CAR-^CTRL^-M did not uptake tumor cells, CAR-M followed two distinct outcomes following engagement with tumor cells (**Movies S1 and S2**). Only 40% of CAR-M quickly engulfed live non-apoptotic FRET^+^ Eµ-Myc tumor cells while a substantial fraction of CAR-M established prolonged and stable contacts with tumor cells without inducing phagocytosis (**Fig. 3C-E, Movies S1-3**). Interestingly, phagocytosis occurred rapidly after a short median contact time of 6 min following tumor addition, further suggesting that CAR-M are intrinsically “poised” for phagocytosis (**Fig. 3E-F**). Similar observations were made with the CD19^+^ MC38 model suggesting that this phenotype is occurring with other tumor types (**Fig. S10, Movie S3**). Importantly, these CAR-M could maintain stable contacts with tumor cells for up to 280 min without subsequent engulfment, defining a population of “non-phagocytic CAR-M” (**Fig. 3G-H and Movie S4**). Thus, these results reveal substantial functional heterogeneity within CAR-M cells, with individual CAR-M adopting either phagocytic and non-phagocytic behaviors upon tumor encounter. This phenotype was not solely attributable to the level of CAR expression as more than 95% of all CAR-M expressed CAR MFI as high as the phagocytic population (YFP^low^), a population only representing 35-40% of all CAR-M (**Fig.S11A**). Moreover, level expression of the don’t- eat-me SIRPα signal was largely similar across phagocytic and non-phagocytic CAR-M populations, underlining the potential of phagocytic determinants not linked to phagocytosis checkpoint expression (**Fig.S11B)**. Instead, phagocytic CAR-M exhibited a larger cellular area, as an indicator of their size and possible membrane availability, compared to non-phagocytic CAR-M (**Fig.S11C**), suggesting that physical parameters may influence the capacity of CAR-M in performing phagocytosis.

**Figure 3.**
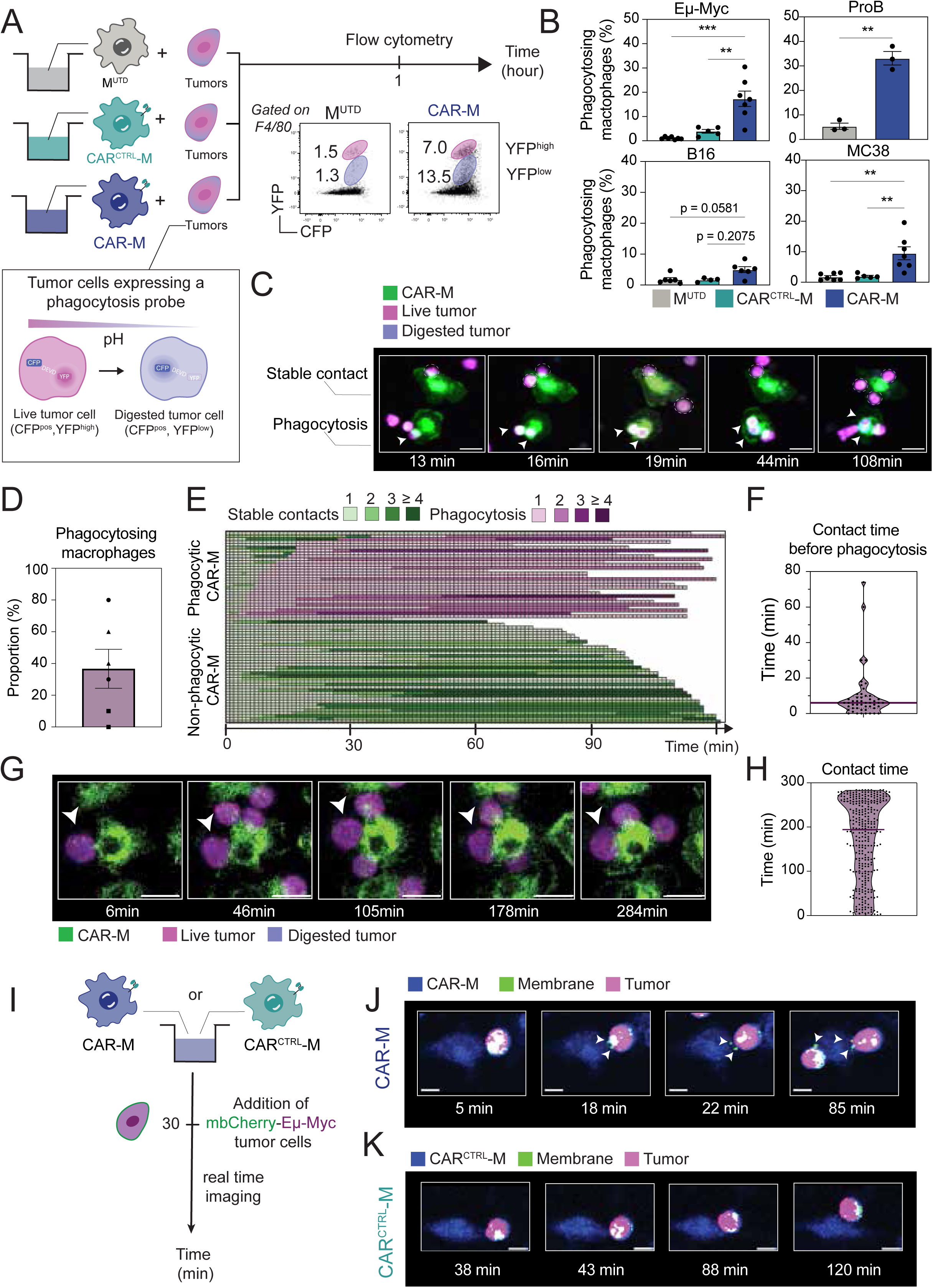
CAR-M tumor interactions result in phagocytic and non-phagocytic outcomes. **A-B**. Experimental scheme (A) of a 1h co-culture of CAR-M with CD19-expressing tumor cells expressing a probe for phagocytosis. Schematic representation of tumor cells expressing the fluorescence phagocytosis probe. At a physiological level, tumor cells express both the CFP and YFP fluorescence (magenta). Upon phagocytosis, YFP fluorescence is gradually lost as tumor cells are engulfed into the highly acidic environment of the phagolysosome, appearing blue during live imaging Quantification (B) of phagocytic CAR-M was carried out as the proportion of F4/80^+^ CAR-bearing (CAR^CTRL^-M and CAR-M) or not (M^UTD^) YFP^low^ cells in different tumor models (n = 3-7 independent experiments, statistical analysis by paired one-way ANOVA (MC38, Eµ-Myc), Kruskal-Wallis test (B16) or paired t-test (ProB)). **C**. Representative time-lapse imaging of the dynamics of CAR-M (green) in co-culture with Eµ-Myc tumors expressing the fluorescence probe. Live tumors appear magenta, engulfed tumor cells are blue. Stable contacts are encircling while an arrow points to the phagocytosis (representative of 3 independent experiments; scale bar, 20 µm). **D.** Proportion of phagocytic CAR-M across all analyzed movies and imaging positions. Each point represents one analyzed position; dot shapes indicate independent experiments (n = 3 independent experiments). **E.** Movie analysis showing interaction dynamics between CAR-M with Eµ-Myc tumor cells during 2 hours of co-culture. Each line represents a CAR-M in contact with a least one tumor cell and each square a minute of acquisition. CAR-M in contact with tumor cells appear green, a phagocytosis event corresponds to a pink square (data pooled from 3 independent experiments, 67 CAR-M analyzed). **F.** Duration of tumor cell contact time preceding phagocytosis events (data pooled from 3 independent experiments, 49 phagocytosis events analyzed). **G**. Representative time-lapse imaging of the dynamics of CAR-M (green) in co-culture with Eµ-Myc tumors expressing the fluorescence probe for 4,5 hours. The arrow indicates a stable contact. Live tumors appear magenta, engulfed tumor blue (representative of 3 independent experiments; scale bar, 20 µm). **H**. Quantification of the duration of contacts between CAR-M and Eµ- Myc tumor cells in time-lapse imaging over 284 minutes (data pooled from 3 independent experiments, 312 stable contacts were analyzed). **I.** Experimental scheme. Briefly, CAR-M or CAR^CTRL^-M were imaged for 30 min before the addition of DEVD-Eµ-Myc tumor cells expressing a membrane targeted mCherry. Dynamic imaging continued for 2 h after. **J.** Representative time-lapse images of the interaction dynamics between CAR-M (blue) and DEVD-mbCherry-Eµ-Myc tumor cells (magenta with green membrane). Time indicates elapsed since tumor addition. Images (representatives of 3 independent experiments) were acquired every minute (scale bar, 10 µm). **K.** Representative time-lapse images of the interaction dynamics between CAR^CTRL^-M (blue) and DEVD-mbCherry-Eµ-Myc tumor cells (magenta with green membrane). Time indicates elapsed since tumor addition. Images were acquired every minute (scale bar, 10 µm) (representative of 3 independent experiments). Error bars represent mean ± SEM. ***P < 0.001; **P < 0.01; *P < 0.05. See Methods for additional experimental details.

To evaluate whether these lasting non-phagocytic CAR-M were totally inert or in contrary still active, we decided to investigate trogocytosis in our system, an active mechanism of membrane exchanges. To this end, tumor cells were transduced to express a membrane-bound mCherry-fluorescent protein allowing visualization of tumor membrane exchanges during CAR-M-tumor cell interactions (**Fig.3I and movies S5-6**). Under these conditions, non-phagocytic CAR-M could be seen nibbling tumor cells and becoming mCherry overtime, an observation largely absent in CAR^CTRL^-M (**Fig.3J-K, movies S5-6**). This suggested that non-phagocytic CAR-M can perform trogocytosis, and that CAR downstream signaling plays an important role in this process. Altogether, these results indicate that CAR-M have both intrinsic phagocytic and non-phagocytic capacities upon tumor interaction which cannot solely be explained by different CAR levels but may relate to intrinsic cellular properties such as cell size and membrane availability. We also show that non-phagocytic interactions can result in trogocytosis events occurring in the presence of a signaling CAR and thereby placing downstream CAR signaling as possibly active even in the absence of phagocytosis.

### Tumor engagement by non-phagocytic CAR-M triggers intracellular calcium signaling

The coexistence of phagocytic and non-phagocytic CAR-M-tumor interactions raises the possibility that sustained CAR-M-tumor interactions that do not result in phagocytic uptake may still be efficient to drive CAR early signaling. Thus, to test this possibility, we analyzed calcium (Ca²⁺) flux, an early signaling event associated with macrophage activation during phagocytosis which we used here as a readout of CAR signaling (24). CAR-M and control macrophages were engineered to express the FRET–based Ca²⁺ sensor Twitch2B (CAR-M^Twitch2B^ , CAR-CTRL^Twitch2B^ Empty^Twitch2B^, respectively), allowing real-time monitoring of intracellular Ca²⁺ dynamics by time-lapse imaging (**Fig. 4A**) (25). As expected, phagocytic CAR-M^Twitch2B^ exhibited one or multiple transient Ca²⁺ fluxes upon tumor contact, followed by rapid target engulfment (**Fig. 4B and Movie S7**). Notably, Ca²⁺ signals were also observed in non-phagocytic CAR- M^Twitch2B^ establishing stable contacts with tumor cells (**Fig.4B-C**). By contrast, control Empty^Twitch2B^ or CAR-CTRL^Twitch2B^ failed to exhibit detectable Ca²^+^ fluxes underscoring the requirement for expressing a full signaling CAR for signal induction (**Fig. 4B-C and Movies S7-S9 and S12**). Together, these results demonstrate that CAR-M–tumor cell interactions induce CAR-dependent Ca²⁺ signaling irrespective of phagocytosis occurrence, providing direct evidence that non-phagocytic CAR engagement initiates intracellular signaling events that may contribute to CAR-M effector functions.

**Figure 4.**
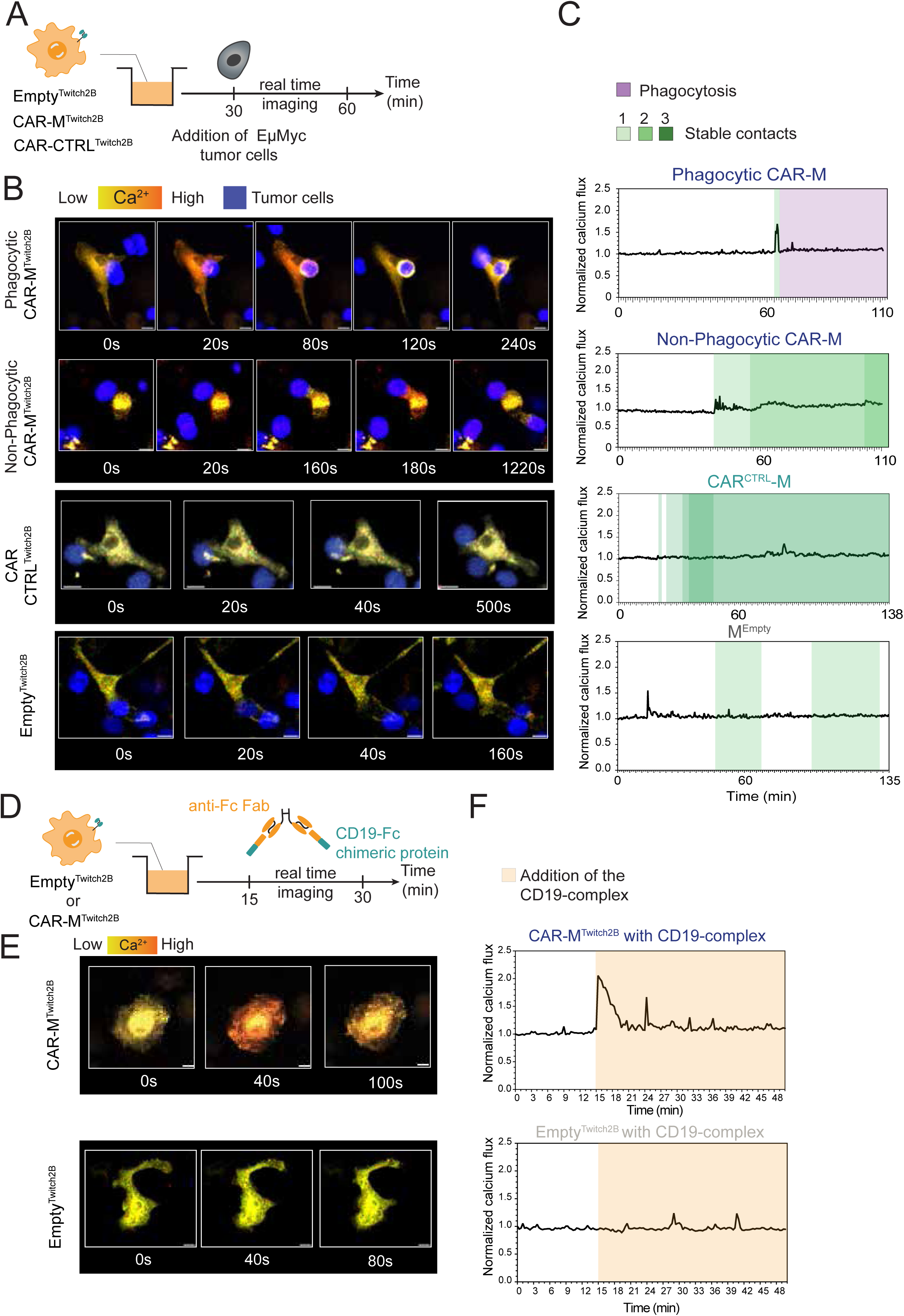
Regardless of outcome, CAR-tumor interactions can trigger calcium signals. **>A-C.** Experimental set up (A) and representative images (B) of a co culture of tumor cells and CAR-M (top) or control macrophages (CAR^CTRL^-M or M^Empty^) expressing (bottom) a calcic probe Twitch2B (CAR- M^Twitch2B^, CAR-CTRL^Twitch2B^ or Empty^Twitch2B^). Images were acquired every 20 s (scale bar, 10 µm). Added Eµ-Myc tumor cells appear blue (representative of 3 independent experiments). Calcium signals were collected from image analyses and are shown in (C). Prolonged contacts with tumor cells appear in green on the quantification, and phagocytosis events in magenta. **D-F.** Experimental scheme (D) and representative images (E) showing live imaging of Twitch 2B-expressing CAR-M or Twitch-2B macrophages (Twitch2B^Empty^) during addition of the CD19-crosslinking complex. Images are representative of 3 independent experiments and were acquired every 20 s (scale bar, 10 µm). (F) Quantification of the calcium signal in CAR-M^Twitch2B^ or Empty^Twitch2B^ macrophages in the presence of the CD19 complex (orange).

Although a significant fraction of CAR-M fails to phagocytose tumor cells, whether these non-phagocytic interactions contribute to the overall CAR-M efficacy remains unclear. To dissociate the impact of CAR signaling from that of phagocytosis, we developed an experimental approach to induce CAR engagement independently of target engulfment. CAR-M^Twitch2B^ were imaged in real time while CAR molecules were crosslinked using a CD19-based complex consisting of a CD19-Fc chimeric protein and a secondary anti-Fc Fab fragment (**Fig. 4D**). Under these conditions, CAR-M^Twitch2B^ displayed a transient strong Ca²⁺ flux that was absent in control Empty^Twitch2B^. Interestingly, moderated Ca^2+^ signaling was recorded in CAR-CTRL^Twitch2B^ lacking a full signaling CAR suggesting that aggregation of CAR molecules at the cell surface is sufficient to trigger some basal level of Ca²⁺ signaling (**Fig. 4E-F, Fig. S12A-C and Movies S10-11 and S13**). These data show that CD19-mediated CAR crosslinking using CD19-Fc chimeric protein can mimic Ca^2+^ signals observed in non-phagocytic CAR-M, therefore providing a tool to model non-phagocytic CAR engagement.

Altogether, these results show that tumor interaction in the absence of phagocytosis can trigger intracellular signaling in CAR-M and occurs only when expressing a full signaling CAR.

### CAR signaling, regardless of target engulfment, is sufficient to induce a pro-inflammatory CAR- M phenotype

We next investigated whether non-phagocytic CAR-M signaling is sufficient to induce CAR-M effector programs. To address this question, we performed an unbiased bulk RNA sequencing of phagocytic CAR-M or CAR-M stimulated by receptor crosslinking, thereby triggering CAR engagement independently of tumor engulfment. Transcriptional profiles were analyzed shortly after CAR engagement and compared to unstimulated CAR-M (**Fig. 5A**). To avoid contamination by engulfed tumor-derived RNA, CAR-M were co-cultured with human Daudi tumor cells engineered to express mouse CD19 and the phagocytosis reporter. This species-mismatched system enabled the selective exclusion of reads mapping to the human genome prior to analysis of macrophage-derived transcripts mapping to the mouse genome, thereby ensuring accurate characterization of CAR-M transcriptional responses (**Fig. S13A**). Given that target engulfment induces rapid (< 1 h) transcriptional reprogramming in macrophages (26), all CAR-M were flow-sorted 1 h after stimulation (**Fig. 5A**). As expected, phagocytosis induced extensive transcriptional remodeling at this early time point (**Fig. S13B**). Strikingly, CAR signaling in the absence of phagocytosis triggered a transcriptional program that largely overlapped with that observed in phagocytic CAR-M (**Fig. 5B-C** and **S13C-D**). Early CAR engagement led to the upregulation of NFκB–associated pathways, as well as cytokine and chemokine transcripts, including signatures related to TNFα and inflammasome activation (**Fig. 5C**). These results indicate that CAR signaling, independently of target engulfment, is sufficient to rapidly induce a pro-inflammatory transcriptional program in CAR-M.

**Figure 5.**
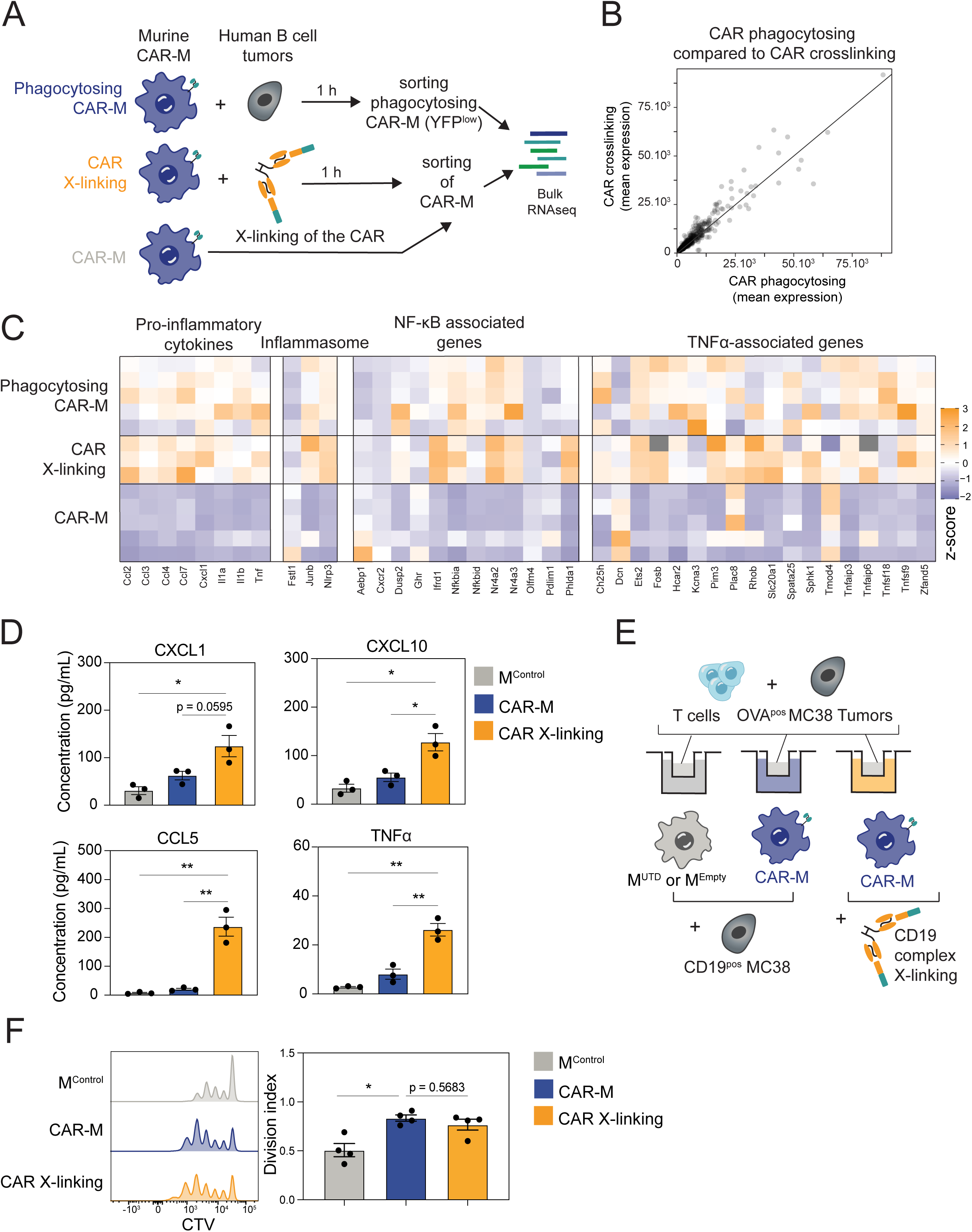
CAR signaling is sufficient for CAR-M mediated-cytokine/chemokine production. **A**. Experimental scheme of the mixed-species bulk RNAseq set-up. Murine purified CAR-M were incubated for 1 h with the CD19-complex, or with human B cell tumors (Daudi) transduced to express murine CD19 and the phagocytosis probe. Control CAR-M did not receive any treatment. Phagocytic CAR-M were identified based on their YFP^low^ fluorescence and flow cytometry sorted. All three conditions were sequenced together. Analysis was performed by first mapping the reads onto the human genome and excluding all reads mapping there, then performing the mapping on the murine genome. The experiment included 3-5 biological replicates and was performed independently twice. **B.** Scatterplot representing the mean expression of each differentially expressed gene (compared to the CAR-M condition) in the CAR phagocytosing condition (x-axis) compared to the CAR crosslinking condition (y-axis). **C**. Heatmap representing the relative expression of genes associated to pro-inflammatory pathways that are significantly differentially regulated in phagocytic or crosslinked CAR- M (n = 3-5 independent experiments). **D.** Absolute concentration of CXCL1, CXCL10, CCL5 and TNFα measured in co-culture of control macrophages (M^Empty^) or CAR-M with Eµ-Myc tumor cells for 24 h. CAR-M were also cultured in the presence of the CD19 complex for 24 h (CAR crosslinking) (n= 3 independent replicates, statistical analysis was performed by paired one-way ANOVA). **E-F**. Schematic experimental set-up (E) showing the transwell assay. Macrophages (Control (M^UTD^ or M^Empty^) or CAR- M) and tumor cells were plated at the bottom part while naïve CTV-stained OT-I T cells were added with MC38-OVA on the upper part of the well. For the crosslinking condition, CAR-M were left without tumor cells at the bottom and the CD19 complex was added. 72 h later T cell proliferation was measured by CTV dilution by flow cytometry and graphed in (F) (n = 4 independent experiments, statistical analysis was performed by Friedman test). Error bars represent mean ± SEM. ***P < 0.001; **P < 0.01; *P < 0.05. See Methods for additional experimental details.

We next asked whether these early transcriptional changes were translated into functional effector responses. To this end, we performed multiplex ELISA analyses to quantify cytokines and chemokines produced by CAR-M cultured either in contact with CD19^+^MC38 tumor cells or stimulated with the CD19 crosslinking complex. TNFα and CXCL1 were also detected in the supernatants under both conditions, along with additional chemokines including CCL5 and CXCL10 (**Fig. 5D**). Notably, concentrations of these mediators were 1.5 times higher with CD19-crosslinking rather than with CAR- M co-cultured with tumor cells (**Fig. 5D**).

To investigate how strength of CAR cross-linking correlates with qualitative or quantitative cytokine production, we cultured CAR-M with different concentrations of CD19 crosslinking complex and performed multiplex ELISA on supernatants, which we compared with CAR-M co-cultured with Eµ- Myc tumor cells (**Fig. S14A**). Cytokine production correlated with increasing amounts of CD19 complex at low doses, although distinct mediators displayed different dynamic responses. Whereas TNFα reached a plateau, chemokines such as CCL5 or CXCL1 continued to increase across a broader range of stimulation intensities possibly reaching a plateau at later stages (**Fig.S14B**). Thus, strength of CAR signaling is likely to drive different profiles of cytokines and chemokines in CAR-M. Overall, these results show that CAR signaling alone, in the absence of target engulfment, is sufficient to drive robust production of pro-inflammatory mediators, establishing phagocytosis-independent CAR activation as an effector mechanism of CAR-M.

Finally, we investigated whether soluble factors produced upon CAR crosslinking were sufficient to enhance T cell activation. Using a transwell system, CTV-labelled naïve OT-I cells were co- cultured with tumor cells and physically separated from CAR-M either engaging tumor cells or stimulated with the CD19 complex (**Fig. 5E**). Under these conditions, CAR crosslinking alone was sufficient to enhance naïve T cell activation, to a degree comparable to that observed with phagocytosing CAR-M (**Fig. 5F**). Collectively, these data demonstrate that CAR signaling, independently of phagocytosis, is sufficient to induce a pro-inflammatory CAR-M phenotype and promote T cell activation, establishing non-phagocytic CAR engagement as a functional and therapeutically relevant mode of CAR-M action.

## DISCUSSION

CAR-M therapy has largely been viewed through the paradigm that tumor cell phagocytosis is the initiating event that triggers all downstream effector functions, including inflammatory cytokine and chemokine production, remodeling of the tumor microenvironment, antigen cross-presentation, and ultimately the activation of T cell-mediated antitumor responses (15). As a result, phagocytic capacity has become the primary readout of therapeutic CAR-M efficacy *in vitro*. However, directly testing whether phagocytosis is required for this assumption has been challenging because clinically relevant CAR-M are generated through adenoviral transduction, a process that imposes a highly inflammatory macrophage state capable of enhancing effector functions independently of CAR engagement. By using a non-polarizing transduction strategy, we were able to allow the specific attribution of CAR-M effector functions to CAR engagement and directly interrogate the contribution of phagocytosis to CAR-M mechanism of action. Unexpectedly, we found that a substantial fraction of CAR-M failed to uptake tumor cells yet remained functionally active. These non-phagocytic CAR-M established sustained contacts with tumor cells that were sufficient to induce trogocytosis, sustained CAR-dependent Ca²⁺ signaling, and a robust pro-inflammatory program. Despite the absence of phagocytosis, CAR-M efficiently promoted T cell activation and proliferation through cytokine and chemokine production, whereas antigen cross-presentation appeared largely dispensable. Together, our findings demonstrate CAR engagement alone, independent of tumor cell engulfment, is sufficient to initiate key immune stimulatory functions of CAR-M, establishing contact-dependent CAR signaling as a previously unrecognized effector mechanism and broadening the current paradigm of CAR-M function beyond tumor cell phagocytosis.

Our findings further suggest that the balance between phagocytic and non-phagocytic outcomes may be shaped by at least two layers of heterogeneity: some that are intrinsic to tumor cells, and others to the macrophages. At the tumor level, the differential phagocytic rates observed across tumor models are consistent with the role of target size in modulating phagocytosis. Although these parameters were not directly quantified here, they are likely to contribute to our observed differences as B cell malignancies harbor a much lower size than mesenchymal tumor targets. Another parameter could be the antigen expression levels and density. Indeed, while absolute expression was quite similar between tumor models, density will inversely correlate to size. While there seems to be a correlation in our hands between these parameters (size and antigen density) and interaction outcomes, we are basing our study on the model antigen CD19. It would be crucial to extend them to a more clinically relevant antigen, such as HER2. Interestingly, similar tumor-intrinsic heterogeneity is likely to exist in the clinical setting, where tumor cells vary widely across cancer types and patients (27) : in the context of “solid” tumors which can harbor very elongated shapes, the phagocytic outcome of CAR-M -tumor interactions thus seems less likely. At the CAR-M level, the capacity to engulf tumor cells is a pre-existing, CAR-independent feature that intrinsically varies within the macrophage population. This heterogeneity could be linked to the origin of the cell population in our set-up. Lineage-related heterogeneity among bone marrow-derived progenitors has been shown to give rise to macrophage subsets with distinct intrinsic phagocytic competencies (28). However, it could also result from biophysical properties such as cell size and membrane availability, consistent with the enhanced phagocytosis observed in myeloid cells lacking G protein subunit Gβ4 exhibited profound plasma membrane expansion (29).

Tumor recognition in the absence of phagocytosis is not a passive or “inert” interaction but instead triggered downstream Ca²⁺ signaling associated with trogocytosis without apparent tumor cell death. These findings demonstrate that productive CAR signaling does not require complete target internalization and suggest that, in this context, trogocytosis primarily reflects CAR signalling rather than serving as a direct cytotoxic mechanism, in contrast to previous reports (30–32). Whether trogocytosis promotes antigen loss and contributes to tumor immune escape, as described for CAR-T cells (33), remains an important question with potential clinical implications. A central question raised by our findings is how non-phagocytic CAR-M, which do not internalize and process tumor antigens, nevertheless promote T cell responses. Our data indicate that antigen cross-presentation is unlikely to represent the primary mechanism in our model. Consistent with this interpretation, previous studies have shown that MHC class I-deficient CAR-M retained partial antitumor activity *in vivo* (20), suggesting that CAR-M-driven adaptive immune activation can occur independently of conventional antigen presentation. Of note, differences in the inflammatory state of CAR-M across studies, driven by distinct transduction strategies, may contribute to the divergent observations reported regarding the contribution of cross-presentation. An alternative contact-dependent mechanism is cross-dressing, whereby macrophages acquire intact MHC-peptide complexes from target cells through membrane transfer, enabling T cell activation without intracellular antigen processing (34). Cross-dressing can be mechanistically linked to trogocytosis, as both involve partial membrane exchange at the immunological synapse. Indeed, it has been shown recently that trogocytosing CAR-M can deplete MHC-I molecules from the target tumor cell surface (30). However, whether CAR-mediated membrane transfer can generate functional peptide-MHC complexes represents an important avenue for future investigation.

Our data support a model in which CAR-dependent cytokine and chemokine secretion constitutes the primary mechanism linking CAR engagement to T cell activation and proliferation. Remarkably, this inflammatory program was initiated independently of phagocytosis, demonstrating that productive CAR signaling alone is sufficient to orchestrate downstream immune responses. Rather than functioning solely as engineered phagocytes, CAR-M therefore emerge as localized inflammatory hubs capable of coordinating the recruitment and activation of multiple immune populations. Among the cytokines upregulated upon CAR engagement, IL-12 is of particular interest because it drives granzyme B expression and cytotoxic capacity in both T and NK cells co-cultured with CAR-M (35). This T cell– macrophage crosstalk acquires particular relevance for solid tumors, where the immunosuppressive tumor microenvironment represents the principal barrier to effective antitumor immunity and a major challenge for cell-based therapies (e.g. CAR T cells) (36). Tumor-associated macrophages are among the most abundant stromal cell types and actively remodel the TME by secreting anti-inflammatory mediators such as IL-10 and TGF-β, while limiting cytotoxic T cell infiltration and activation (2, 36). In this context, the capacity of CAR-M to produce a pro-inflammatory, independently of phagocytosis, positions them as potent remodelers of the immunosuppressive solid tumor TME. By locally releasing cytokines and chemokines including IL-12, TNFα, CXCL10, and CCL5, CAR-M may simultaneously reactivate exhausted tumor-infiltrating T lymphocytes, promote their recruitment at the tumor site, and reprogram bystander immunosuppressive TAMs toward a pro-inflammatory state, thereby amplifying the global antitumor immune loop. Such amplification of local immune circuits could overcome T cell exclusion and exhaustion, thereby enhance tumor elimination (37–39). These findings suggest that optimizing the inflammatory “secretome” of CAR-M may represent a more efficient strategy for solid tumor than maximizing phagocytic efficiency alone.

Our findings have important implications for the design of next-generation CAR-M therapies and strengthen the rationale for incorporating signaling domains that engage pro-inflammatory cytokine pathways, rather than focusing exclusively on enhancing phagocytic capacity. Consistent with this concept, recent studies have shown that third-generation CARs activating TLR4 intracellular signaling can induce tumor cell apoptosis and CAR-M-mediated efferocytosis, leading to enhanced survival in preclinical models (40). Extending such engineering strategies to other cytokine pathways, including IL- 12 or TNFα signaling axes, represents a promising direction for future work. Importantly, cytokine-producing CAR-M may also enable a bystander mode of action, facilitating the elimination of antigen-negative tumor cells that would otherwise escape phagocytosis-dependent killing. An important underexplored question concerns the durability of CAR-M inflammatory program. Sustained CAR engagement, particularly in non-phagocytic contexts, may expose CAR-M to prolonged stimulation without resolution through target engulfment, potentially leading to a state analogous to T cell exhaustion and progressive loss of cytokine production. Determining whether non-phagocytic CAR-M are susceptible to exhaustion, and whether specific intracellular signaling domains can prevent it, will be essential to fully exploit the cytokine-driven mechanisms identified here.

Ultimately, this study establishes CAR engagement, regardless of target engulfment, as a key contributor to CAR-M overall mode of action. Our work therefore broadens the current view of CAR-M biology by identifying cytokine-driven immune activation as a major component of their therapeutic activity and opens new avenues for the rational design of next-generation CAR-M therapies.

## MATERIAL AND METHODS

### Mice

Five-to-twelve-week-old female and male C57BL/6J were purchased from Envigo. *Rag2*-deficient mice, Ubi^GFP^-OT-I, OT-I *Rag2*^−/−^ and Ly5.1^+^*Rag2*^−/−^ under a C57BL/6 background were bred in our animal facility under specific-pathogen-free conditions, air renewal, ambient temperature of 22 ± 2 °C and controlled light–dark cycle. All experiments were approved by Institut Pasteur’s Safety Committee in accordance with French and European guidelines (CETEA 220108, CETEA 240110, CETEA 230038).

### Plasmids

All used plasmids were subcloned into a MSCV backbone. The CAR plasmid encodes a chimeric antigen receptor composed of an extracellular single chain variable fragment (scFv) targeting mouse CD19, a CD8 transmembrane domain, and an intracellular domain containing the FcR-γ immunoreceptor tyrosine-based activation motif (ITAM) and the p85-recruiting motif of murine CD19 (17). The CAR^CTRL^ plasmid encodes a chimeric antigen receptor only comprising the extracellular scFv targeting mouse CD19 and transmembrane domain, lacking any intracellular signaling motives (41). Furthermore, plasmids encoding the probes DEVD (42) and Twitch2B (25), gifted by Oliver Griesbeck, were used. The cytoplasmic ovalbumin (OVA) encoding plasmid was subcloned from pCI-neo-cOVA, a gift from Maria Castro (Addgene plasmid # 25097 ; http://n2t.net/addgene:25097 ; RRID:Addgene_25097) (43). The membrane targeted mCherry plasmid sequence was a gift from Catherine Berlot (Addgene plasmid # 55779 ; http://n2t.net/addgene:55779 ; RRID:Addgene_55779).

### Cell lines

The leukemic ProB cell line (44) and the lymphoma myc-over expressing cell line (Eµ-myc mice (45)) were retrovirally transduced to express a fluorescent-based phagocytosis reporter probe (named here DEVD) (23). Human B cell Daudi tumors (ATCC, CCL-213) were retrovirally transduced to express the DEVD phagocytosis reporter as well as murine CD19 and CD20 molecules. All B cell tumor models were cultured in RPMI-1640 (Gibco) supplemented with 10% heat-inactivated fetal bovine serum (Sigma), 50 U.mL^-1^ penicillin (Gibco), 50 μg.mL^-1^ streptomycin (Gibco), 1 mM sodium pyruvate (Gibco), 10 mM HEPES (Gibco), and 50 μM 2-mercaptoethanol (Gibco) (complete RPMI). The melanoma cell line B16_F10 (referred to as B16, ATCC) was retrovirally transduced to express the DEVD probe as well as the murine CD19 protein (46). HEK293T (ATCC) and L929 (ATCC) and the generated B16-DEVD-CD19 were cultured in DMEM (Gibco) supplemented with 10% heat-inactivated fetal bovine serum (Sigma), 50 U.mL^-1^ penicillin (Gibco), 50 μg.mL^-1^ streptomycin (Gibco), 1 mM sodium pyruvate (Gibco), 10 mM HEPES (Gibco), and 50 μM 2-mercaptoethanol (Gibco) (complete DMEM). The colon carcinoma cell line MC38 (Kerafast) was modified to express the DEVD probe as well as murine CD19 protein (46). When indicated, the cells were further genetically modified to express the cytosolic ovalbumin protein. Cells were cultured in DMEM (Gibco) supplemented with supplemented with 10% heat-inactivated fetal bovine serum (Sigma), 50 U.mL^-1^ penicillin (Gibco), 50 μg.mL^-1^ streptomycin (Gibco), 1 mM sodium pyruvate (Gibco), 10 mM HEPES (Gibco), 50 µM gentamicin (Sigma) and 1mM non-essential amino acids (Sigma). All cell lines were cultured at 37°C, 5% CO2 and regularly tested for mycoplasma contamination (Minerva Biolabs).

### Generation of L-cell conditioned medium (L929 supernatant)

L929 cells were plated in 175 mL cell-culture treated flasks in the above-described DMEM-based medium. Once the cells reached confluence, the culture medium was removed and replaced with 100 mL of fresh medium. 7 days later, the medium was collected, filtered on 0.2 µm filters and aliquoted for storage at −20°C.

### Generation of β2m-deficient tumor cell lines

DEVD^+^ CD19^+^ OVA^+^ β2m-deficient MC38 tumor cells were generated from the parental line MC38 DEVD^+^ CD19^+^OVA ^+^ using the genotype editing Alt-RTM CRISPR-Cas9 System (Integrated DNA Technologies) and following manufacturer’s recommendations. The sequences used to target β2m was CACGCCACCCACCGGAGAAT. Effective β2m ablation was confirmed by flow cytometry.

### Virus generation for bone-marrow transduction

HEK293 cells were plated in 10 cm^2^ dishes. Once they reached 70-90% of confluence, the cells were transfected with 2 µg of pCL-Eco vector and 3 µg of plasmid of interest using JetPRIME® transfection reagent (Polyplus transfection) according to the manufacturer’s guidelines. 4-6 h later, the cell culture medium was replaced with 7 mL complete RPMI. 24h and 48h after transfection, the medium was collected and filtered on 0.45 µm filters.

### Generation of bone-marrow derived macrophages (BMDMs)

Bone-marrow derived macrophages were generated as described previously (47). Briefly, bone-marrow was extracted from C57BL/6J and cultured for at least 6 days with 20% L929 supernatant in complete RPMI. Transduced macrophages were generated by two rounds of spin-infections (at 4 and 24 h post-extraction) performed at 32°C for 1,5 h and 1000g with retroviral particles generated as described above and supplemented with 10 µg mL^-1^ of polybrene (Merck). In case of double transduction, viral particles encoding each one plasmid were mixed at a 1:1 ratio. After the second spin-infection, the cells were detached and plated in non-tissue-culture treated 15 cm dishes with complete medium supplemented with 20% L929 supernatant. Once the cells were differentiated, the macrophages were detached using cold 1X PBS followed by a 15 min incubation with cell dissociation buffer (PBS supplemented with 5 mM EDTA) at 4°C.The transduction rate (around 50-80% of the cells) was screened by flow cytometry using cell surface expression of human CD34 reporter. Unless specified otherwise, macrophages were always kept in complete medium supplemented with 20% L929 supernatant.

### Purification of CAR-M

For Luminex, dynamic imaging and transwell or co-culture with T cells CAR^CTRL^-M or CAR-M were purified according to their CAR expression as defined by cell surface expression of human CD34 protein using hCD34-positive selection following the manufacturer’s instructions, either by directly using anti-hCD34 beads or using hCD34-PE staining followed by a separation using anti-PE beads (Miltenyi). Purification efficiency (>70%) was screened by flow cytometry.

### CD19 crosslinking complex

To generate the CD19 complex, the murine CD19-Fc chimera (R&D Biosystems) was incubated with anti-Fc Fab (Jackson Immunoresearch Laboratories) at a ratio of 1:2 for at least 10 min before addition to the cells for a final concentration of 2.5 µg.mL^-1^ of CD19-Fc and 5 µg.mL^-1^ anti-Fc Fab.

***In vitro* phagocytosis analysis.** Untransduced BMDM, CAR^CTRL^-M or CAR-M were plated in non-tissue culture treated wells. On the day of the assay, tumor cells (Eµ-Myc, ProB, MC38 or B16) were added to the macrophages at an effector:target ratio of 1:1 for 1 h. The supernatant of the wells was removed, the cells washed with cold PBS and incubated for 15 min at 4°C with cell dissociation buffer (Gibco) to allow detachment by pipette flushing.

### *In vitro* antitumoral activity analysis

Untransduced BMDM, CAR^CTRL^-M or CAR-M were plated in non-tissue culture treated wells. On the day of the assay, tumor cells (Eµ-Myc, ProB, MC38 or B16) were added to the macrophages at an effector: target ratio of 1:1 for 24 or 48 h. The supernatant of the wells was removed, the cells washed with cold PBS and incubated for 15 min at 4°C with cell dissociation buffer (Gibco) to allow detachment by pipette flushing.

### Real-time phagocytosis dynamic imaging

Untransduced macrophages, CAR^CTRL^-M or CAR-M were plated at a concentration of 10^6^ cells/well in 6 cm dishes for at least 24 h before imaging. Imaging was performed as described previously (47). Briefly, on the day of imaging, the macrophages were stained using anti-F4/80-Alexa594 (clone BM8, Biolegend) in PBS supplemented with 0.05% FCS for 20 min at room temperature in the dark. The staining solution was removed and replaced with imaging medium (RPMI lacking phenol red (Gibco) supplemented with 5% FCS). Macrophages were imaged at steady-state before addition of 10^6^ cells Eµ-Myc DEVD^+^ or MC38 DEVD^+^CD19^+^ tumors. Acquisition was then performed for at least 2 h. *In vitro* imaging was performed with an upright microscope (FVMPE-RS, Olympus), a 25×/1.05 numerical aperture, water-dipping objective combined with an objective heater and using FV31S-SW software (Olympus). Excitation was provided by an Insight DeepSee dual laser (Spectra-Physics) tuned at 830 nm. The following filters were used for fluorescence detection: DEVD (CFP:483/32, FRET: 542/27), A594 (593/40). Acquisition was also performed on the upright Stellaris Dive microscope (Leica), a 25×/1.05 numerical aperture, water-dipping objective combined with an objective heater (Olympus) and using the Leica Application Suite X (v 4.5.0.25531, Leica). Excitation was provided by a Coherent Vision II laser (BD Biosciences) tuned at 830 nm. The following detection settings were used: CFP (449-505 nm), FRET (530-550 nm), A594 (560-600 nm) and background (620- 650 nm). To create time-lapse sequences, we typically scanned a 20 μm-thick volume at 4 μm Z-steps and 1 min intervals.

### Real-time trogocytosis dynamic imaging

CAR^CTRL^-M or CAR-M were plated at a concentration of 10^6^/well cells in 6 cm dishes for at least 24 h before imaging. Imaging was performed as described previously (47). Briefly, on the day of imaging, the macrophages were stained using anti-F4/80- Alexa488 (clone BM8, Biolegend) and 5 µM CFSE (Invitrogen) in PBS for 20 min at 37°C in the dark. The staining solution was removed and replaced with imaging medium (RPMI lacking phenol red (Gibco) supplemented with 5% FCS). Macrophages were imaged at steady-state before addition of 10^6^ cells Eµ-Myc DEVD^+^ mbCherry^+^. Acquisition was then performed for at least 2 h. *In vitro* imaging was performed with an upright microscope (FVMPE-RS, Olympus), a 25×/1.05 numerical aperture, water-dipping objective combined with an objective heater and using FV31S-SW software (Olympus). Excitation was provided by an Insight DeepSee dual laser (Spectra-Physics) tuned at 880 nm. The following filters were used for fluorescence detection: DEVD (FRET: 542/27), GFP (512/25) and mCherry (624/40). To create time-lapse sequences, we typically scanned a 20 μm-thick volume at 4 μm Z-steps and 1 min intervals.

### Real-time imaging of calcium signaling

Bone-marrow derived macrophages transduced with the retrovirus encoding the Twitch2B probe (Empty^Twitch2B^), the non-signalling CAR CTRL or the CAR-encoding retrovirus and the Twitch2B probe (respectively CAR-CTRL^Twitch2B^ or CAR-M^Twitch2B^). Imaging was performed as described above: macrophages were imaged for 30 min at steady-state before the addition of 10^6^ cells Eµ-Myc tumor cells stained with 5µM SNARF dye (Invitrogen). Acquisition was then continued for at least 1h. In parallel, macrophages were imaged for 15-20 min at steady-state before the addition of the complexed CD19 antigen. Acquisition was then continued for at least 30 min. Acquisition was performed on the upright Stellaris Dive microscope (Leica), a 25×/1.05 numerical aperture, water-dipping objective combined with an objective heater (Olympus) and using the Leica Application Suite X (v 4.5.0.25531, Leica). Excitation was provided by a Coherent Vision II laser (BD Biosciences) tuned at 880 nm. The following detection settings were used: SNARF (637-665 nm), SNARF (571-604 nm), calcium high (536-558 nm), and calcium low (468-515 nm). To create time-lapse sequences, we typically scanned a 16 μm-thick volume at 4 μm Z-steps and 20s intervals.

### Real-time imaging analysis

Movies were processed and analyzed using Fiji software (Version 2.14.0/1.54f). Figures and videos based on two-photon microscopy are two-dimensional maximum or average intensity projections of three-dimensional data. CAR-M-tumor interactions were quantified manually. A prolonged interaction is defined as any interaction lasting at least 1 min. Phagocytosis was defined as the moment the tumor cell was fully surrounded by macrophage cell membrane. Size and circularity of the macrophages were quantified using Fiji software (Version 2.14.0/1.54f) and are based on the morphology of the CAR-M before tumor addition. Calcium signaling was quantified using Fiji (2.14.0/1.54f). Briefly, macrophages of interest were contoured, and median fluorescence intensity in the calcium high and calcium low channels were extracted for each frame. The ratio of calcium high/calcium low was then computed and scaled to the first frame of the movie.

### Mixed-species bulk RNAseq analysis

M^Empty^ or CAR-M were plated at 2×10^6^ cells/well in 6-well untreated cell culture plates. On the day of the experiment, either Daudi-DEVD-mCD19-mCD20 tumors were added at an effector:target ratio of 2:1, or the CD19 antigen complex was added before 1 h of incubation. Cells were then detached as previously described and filtered on 70 µm filters. The single-cell suspension was stained in a Fc-blocked flow cytometry buffer (2% FBS, 0.2% EDTA in PBS) containing anti-CD16/32 antibody (BioLegend) for 20 min at 4°C with the monoclonal antibodies: anti-human CD34 Alexa 647 (clone 561, Biolegend) and anti-F4/80 BV711 (clone BM8, Biolegend). Phagocytosing macrophages were sorted by gating on high CAR-expressing macrophages and then on the YFP^low^ population. Sorting was performed on FACS Aria III (BD Biosciences) using a 100 µm nozzle. Cells were collected in FCS supplemented with 1 mM EDTA (Gibco). RNA extraction was then performed using the RNAeasy extraction kit (Qiagen). RNA concentrations were measured on the Qubit (Invitrogen) and a quality control performed on the TapeStation (Agilent). mRNA libraries were generated and sequenced at the internal platform (Biomics) using the Index Dual RNA kit (IDT). Libraries were sequenced on the Illumina NovaSeq 2000 (100bp, single-end, 100M reads per sample). Raw data quality was assessed, and reports generated using FastQC v.0.12.1 and gathered using multiqc v1.25. We performed a 3 steps mapping: sequencing reads were first mapped to the pMSCV- CARmacro-1D3-tandem-hCD34 using BWA-MEM v0.7.19-r1273. The unmapped reads were recovered using samtools v1.21. These reads were mapped to human genome GRCh38 downloaded from ensembl 112 using STAR v2.7.10b. Then the reads not mapped on the human genome were mapped to the mouse genome GRCm39 downloaded from ensembl 112. After mapping, we used feature Counts from Subread (version 2.0.6) to generate the gene-level count matrix. We performed the matrix normalization (median of ratio method) and differential gene expression analyses by the package DESeq2 v 1.50 in R 4.5.0. Differentially expressed genes were defined as genes with a false discovery rate (FDR) ≤ 0.05 and |Log2FC| ≥ 0.584. Plots were generated using R Studio (v2024.12.1+563).

### Multiplex assays for cytokine quantification *in vitro*

M^UTD^, M^Empty^, CAR^CTRL^ or CAR-M macrophages were plated in the presence or absence of Eµ-Myc or MC38 tumors at an effector: target ratio of 1:1 for 24 h. For the crosslinking condition, the CD19-antigen complex was added to CAR-M for 24 h at 37°C at different concentrations including 2,5 µg ml^-1^, 0,5 µg.mL^-1^, 0,25 µg.mL^-1^, 0,05 µg ml^-1^, 0,025 µg ml^-1^. M^UTD^ were also incubated with 1 µm ml^-1^ LPS (Sigma) for 24 h. At the end of the assay, the supernatant was collected from the cells and spun down at 300g for 5 min to remove any cellular debris. Multiplex cytokine assay was performed using a custom 17-Plex ProcartaPlex Panel (Invitrogen). Plates were incubated overnight at 4 °C under agitation and analyses were performed using a Bio-Plex 200 system and the Bio-Plex Manager software (Bio-Rad). For the MC38 condition, data were normalized to the tumor alone condition. Data are represented as a z-score calculated using this formula: 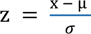 where x is the value of interest, µ the average of all the values of the sample and σ the standard-deviation of all values of the samples.

### CAR-M T cell co-culture assay

Naïve OT-I CD8^+^ T cells were isolated from (Ubi^GFP^)-OT-1 TCR- *Rag1*^−/−^ transgenic mice, labeled with the Cell Trace Violet Proliferation Dye (CTV; Thermo Fisher Scientific) according to the manufacturer’s instructions. Untransduced or CAR-bearing macrophages were co-cultured with MC38-DEVD^+^-CD19^+^-OVA^+^ or MC38-DEVD^+^-CD19^+^-OVA^+^-*β2m*^-/-^ tumors **at an** effector-to-target ratio of 5:1. After 4 h of co-culture, tumor cells were washed by gentle PBS flushing (Gibco). For the OVA-peptide pulsing condition, the SIINFEKL peptide (GeneCust) was added at a concentration of 10 µg ml^-1^ for 4 h before PBS washing. Then CTV-labeled naïve OT-I T cells were added at a T cell to macrophage ratio of 5:1 in the presence of 10 ng ml^-1^ human IL-2 (R&D Systems).

### Transwell assays

A total of 0.5 x10^6^ control macrophages (M^UTD^ or M^Empty^), CAR^CTRL^-M or CAR-M were plated at the bottom of 24-well transwell bearing plates (Corning). On the day of the experiment, MC38-DEVD^+^-CD19^+^-OVA^+^, Eµ-Myc-DEVD^+^ or MC38-DEVD^+^ tumor cells were added to the macrophages at an effector-to-target ratio of 5:1 and incubated for 4h. Tumor cells were washed with PBS. 10^4^ MC38-DEVD^+^-CD19^+^-OVA^+^ or no tumor cells were added to the top of transwell with 0.8 x10^6^ naïve OT-I T cells, generated as described above to mimic the tumor microenvironment and antigen presence. For the cross-linking condition, macrophages were stimulated in the absence of tumor cells using a CD19-antigen complex (2.5 µg ml^-1^ final concentration). All cultures were maintained for 72 h in the presence of human IL-2 (R& D Systems; 10 ng ml^-1^). In the last 4 h, GolgiPlug^TM^ (BD Biosciences) was added to the culture according to manufacturer’s instructions.

*In vivo* i.p. tumour mouse model

C57BL/6 female or male mice of 6-12 weeks were injected i.p. with 4 x10^6^ ProB-DEVD (male tumor), 5 x10^6^ Eµ-Myc-DEVD^+^ (male tumor) or 5 x10^6^ MC38-DEVD^+^-CD19^+^- OVA^+^ (female) tumors respectively 48 or 24 h before treatment with PBS, untransduced, empty or CAR-bearing macrophages differentiated from *Rag2*^−/−^ x Ly5.1 mice bone-marrow as described above using complete RPMI medium supplemented with 20 ng ml^-1^ m-CSF (Peprotech). 1 or 2 days post-treatment, a peritoneal lavage was performed. The collected cells were then spun down and resuspended in red blood cell lysis buffer (Invitrogen) for 5 min. After addition of PBS, the suspension was filtered on 70 µm filters, resuspended and stained.

### Multiplex assays for cytokine quantification *in vivo*

PBS, M^UTD^, or CAR-M treated mice bearing Eµ- Myc were sacrificed 24h after treatment according to ethical standards. Peritoneal lavages were performed and collected fluids spun down for 4 min at 300g. The pellets were used for further flow cytometry analysis. For multiplexed ELISA quantification, supernatants were collected and concentrated using Amicon Ultra-4 centrifugation units (Merck). Multiplex cytokine assay was performed using a custom 17-Plex ProcartaPlex Panel (Invitrogen). Plates were incubated overnight at 4 °C under agitation and analyses were performed using a Bio-Plex 200 system and the Bio-Plex Manager software (Bio-Rad). Data are represented as a z-score calculated using this formula: 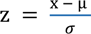 where x is the value of interest, µ the average of all the values of the sample and σ the standard-deviation of all values of the samples.

### Flow cytometry

All flow cytometry stainings were done in flow cytometry buffer (2% FBS, 0.2% EDTA in PBS) with anti-CD16/32 antibody (BioLegend), mice serum (1:50), unless otherwise specified.

### Antibody mix for CAR-M transduction efficacy

Cells were stained with fixable viability dye eF780 (Invitrogen) for 20 min using following monoclonal antibodies: anti-human CD34 A647 (clone 561, Biolegend), anti-human CD34 PE (clone 561, Biolegend), anti-F4/80 BUV395 clone BM8, eBiosciences), anti-Ly6C BV786 (clone HK1.4, Biolegend), anti-G4S Linker A647 (clone E702V, Cell Signaling).

### Antibody mix for phagocytosis and antitumoral activity assays *in vitro*

Cells were stained with fixable viability dye eF780 (Invitrogen) for 20 min using following monoclonal antibodies: anti-IA/IE BUV395 (clone M5/114.15.2, BD Biosciences), anti-CD19 BV650 (clone 6D5, Biolegend), anti-F4/80 BV711 (clone BM8, Biolegend), anti-human CD34 PE (clone 561, Biolegend), anti-CD86 PE-Cy7 (clone GL-1, Biolegend), anti-CD64 APC (clone X54-5/7.1, Biolegend), anti-SIRPα A700 (P84, Biolegend).

### Antibody mix for CAR-M and T cell co-culture

Cells were stained with fixable viability dye eF780 (Invitrogen) for 20 min using following monoclonal antibodies: anti-IA/IE BUV737 (clone M5/114.15.2, BD Biosciences), anti-PDL1 BUV395 (clone MIH5, BD Biosciences), anti-F4/80 PE-Cy7 (clone BM8, Biolegend), anti-CD45.2 BV605 (clone 104, Biolegend), anti-CD40 A647 (clone HM40-3, BD Biosciences), anti-human CD34 PE (clone 561, Biolegend), anti-CD86 BV650 (clone GL-1, Biolegend), anti-TCRβ BUV395 (clone 53-6.7, BD Biosciences), anti-F4/80 BV711 (clone BM8, Biolegend), anti-PD-1 (clone RMP1-14, Biolegend), anti-CD69 APC (clone H1.2F3, Biolegend).

### Antibody mix for transwell assay

Cells were stained with fixable viability dye eF780 (Invitrogen) for 20 min using following monoclonal antibodies: Anti-CD8 BUV395 (53-6.7, BD Biosciences), anti-CD44 BV605 (clone IM7, Biolegend), anti-F480 BV711 (clone BM8, Biolegend), anti-CD25 A488 (clone PC61, Biolegend), anti-PD-1 PE-Cy7 (clone RMP1-14, Biolegend), anti-CD62L APC (clone MEL-14, Biolegend), anti-CD8 PE-Cy7 (clone 53-6.7, Biolegend) and anti-F4/80 APC-Cy7 (clone BM8, Biolegend). For intracellular protein staining, the samples were fixed and stained using the CytoFix CytoPerm kit according to manufacturer’s instruction (BD Biosciences). The following antibodies were used: Anti-IFNγ BUV737 (clone XMG1.2, BD Biosciences), anti-TNFα A488 (clone MP6-XT22, Biolegend), anti-Granzyme B PE (clone QA18A28, Biolegend), anti-Perforin APC (clone S16009B, Biolegend).

### Antibody mix for *in vivo* tumor load analysis

Single cell suspension obtained as described above were stained with fixable viability dye eF780 (Invitrogen) for 20 min using following monoclonal antibodies: Anti-CD19 BUV395 (clone 1D3, BD Biosciences), anti-CD45.2 BUV737 (clone 104, BD Biosciences), anti-B220 BV786 (clone RA3-6B2, Biolegend), anti-NK1.1 PE (clone PK136, Biolegend), anti-human CD34 (clone 561, Biolegend), anti-CD45.1 PE-Cy7 (clone A20, Biolegend), anti-CD8 A647 (clone 53-6.7, Biolegend), anti-TCRβ BV421 (clone 53-6.7, Biolegend), anti-CD45.1 PE (clone A20, Biolegend) anti-F4/80 PE-Cy7 (clone BM8, Biolegend) anti-B220 APC (clone RA3-6B2, Biolegend), anti-human CD34 A700 (clone 581, Biolegend), anti-TCRβ APC-Cy7 (clone 53-6.7, Biolegend), anti- Ly6G BUV395 (clone 1A8, BD Biosciences), anti-Ly6C BV786 (clone HK1.4, Biolegend), anti-CD19 BV421 (clone 6D5, Biolegend), anti-CD11b PerCP5.5 (clone M1/70, Biolegend), anti-CD64 APC (clone X54-5/7.1, Biolegend). For evaluation of absolute counts of cells, counting beads (AccuCheck or 123count, ThermoFisher) were added before acquisition on the LSR Fortessa (BD Biosciences) or Cytoflex (Beckman-Coulter). All flow cytometry data were analyzed using FlowJo v10.

### Proliferation analysis

T cell proliferation was analyzed using the Proliferation analysis tool of Flowjo v10. The division index is defined as the average number of divisions for all cell culture calculated as such: 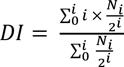 where i is the generation number and N^i^ the number of events in generation i.

### Statistical analysis

Each biological replicates *in vitro* refers to an independent production of macrophages. *In vivo*, each biological replicate corresponds to one mouse. All statistical tests were performed with Prism v.10.6.1 (GraphPad). Data are expressed as mean ± SEM. After normality testing (Shapiro-Wilk test), we used t-tests or non-parametric tests for two-group comparison and one-way or two-way analysis of variance (ANOVA), as well as non-parametric tests (e.g. Kruskal-Wallis’s test) for multiple comparisons. The exact test used for each statistical analysis is indicated in the figure legend. All statistical tests were two-tailed with a significance level of 0.05. *P < 0.05, **P < 0.01, ***P < 0.001; ns, non-significant.

## ACKNOWLEDGMENTS

First and foremost, we would like to dedicate this study to Philippe Bousso who unfortunately passed away before witnessing the publication of this work. We would like to heartfully thank him for supporting us throughout the years with his remarkable scientific guidance, his inspiring rigor and his kind mentorship.

Second, we thank all members of the Bousso laboratory for their support and helpful discussions. Particularly, we thank Nicolas Serafini for critical review of this study. We would like to thank Sébastien Lemoine and Emeline Perthame for their advice about bulk RNAseq analysis. We would also like to thank Ophélie Godon and Friederike Jonsson for sharing reagents and technical input. At the Institut Pasteur, we acknowledge the Central Animal Facility, in particular animal caretakers for their daily help, the CBUTechS platform for cytometry support, and the BIOMICS platform for the Bulk RNA Seq experiment, especially Yakov Vitrenko and Laure Lemée Biomics Platform, C2RT, Institut Pasteur, Paris, France, supported by France Génomique (ANR-10-INBS-09) and IBISA. This work is supported by grants from Institut Pasteur (P.B, L.F.), Institut National de la Santé et de la Recherche Médicale (Inserm, P.B, C.LG), European Research Council (ENLIGHTEN, P.B), La Ligue contre le Cancer (P.B.), ANR JC/JC (CARPHA, C.L.G) and a Ph.D funding from Studienstiftung des deutschen Volkes (Promotionsstipendium, L.F.) and La Fondation ARC (ARCDOC42024120009175, L.F).

## AUTHOR CONTRIBUTIONS

According to the CrediT taxonomy (48). Conceptualization: C.L.G.; Methodology: L.F., A.V., F.L., B.C., Z.G., C.L.G.; Investigation: L.F., C.V., R.A., B.C., C.L.G.; Data curation: L.F., A.V.; Formal analysis: L.F., A.V., Visualization: L.F., C.L.G.; Writing – original draft: L.F., M.V.G., C.L.G.; Writing – review & editing: L.F., F.L., C.V., R.A., B.C., M.V.G., C.L.G.; Resources: P.B.; Supervision: P.B., M.V.G, C.L.G.; Project administration: C.L.G.; Funding acquisition: P.B., C.L.G.

## MATERIALS AND CORRESPONDENCE

All data needed to evaluate the conclusions are present in the paper and the supplementary materials. Further information for the study should be directed to and will be fulfilled by the corresponding author C.L.G., upon reasonable request.

## SUPPLEMENTARY FIGURE LEGEND

**Figure S1. Generation and characterization of CAR-M.**

A. Experimental scheme of the transduction protocol of CAR-M using retroviral infection. **B.** Representative dot plot of the expression of F4/80 and Ly6C by CAR-M after 6 days of differentiation *in vitro*. **C.** Representative histogram of the transduction level of M^UTD^ or CAR-M measured by the level of hCD34 expression after purification of the CAR-M. Representative of more than 10 independent experiments. **D.** Representative contour plot of the expression of the CAR on M^UTD^ (grey) or CAR-M (blue) as measured by hCD34 expression versus a direct CAR-staining using the anti-linker antibody.

**Figure S2. Absolute concentrations in the supernatant of M^UTD^, M^Empty^ or CAR-M at steady-state of different cytokines and chemokines.**

Supernatants were collected at 24 h post-stimulation (n = 3-11 independent experiments, statistical analysis by one-way ANOVA for normally distributed samples (CXCL1, CXCL10, CCL2, CCL4 and CCL5) and by non-parametric Kruskal-Wallis test for non-normally distributed samples (Il-10, IL-6, IL- 12, MIP2α and TNF-α)). Error bars represent mean ± SEM. See Methods for additional experimental details. *\*\*\*P < 0.001; **P < 0.01;* *P < 0.05.

**Figure S3. Antitumoral activity of CAR-M *in vitro* and *in vivo*.**

**A.** CD19 expression on the surface of different tumor models including CD19^neg^ MC38, Eµ-Myc, CD19^+^ B16 and CD19^+^ MC38 (representative of > 8 independent experiments). **B.** Normalized tumor load of CD19^−^ MC38 in the presence of M^UTD^ or CAR-M after 48 h. Tumor load is normalized to the condition of tumor cells alone (n = 5 independent experiments, statistical analysis by paired t-test). **C.** Experimental scheme. **D.** Tumor load 48 h after i.p. injection of CAR-M or control macrophages (M^Empty^) into ProB-bearing C57BL/6 mice. Tumor load was measured by flow cytometry and normalized to the control condition (n = 7 mice from 2 independent experiments; statistical analysis by Mann-Whitney). **E.** Proportion of endogenous peritoneal B cells (CD45.2^pos^, B220^pos^) of ProB-bearing C57BL/6 mice treated with M^Empty^ or CAR-M (n = 7 mice from 2 independent experiments; statistical analysis by Mann-Whitney). Error bars represent mean ± SEM. ***P < 0.001; **P < 0.01; *P < 0.05. See Methods for additional experimental details.

**Figure S4. *β2m^−/−^* tumor controls and T cell proliferation flow cytometry profile.**

**A.** Representative histograms of the expression levels of MHC-I (H2kB staining) of WT, *β2m*-deficient or unstained tumor cells. **B.** Representative histograms of the proliferation profile (CTV) of naïve OT-I T cells in presence of M^UTD^ or CAR-M primed with WT or *β2m*-deficient tumor (n = 3-5 independent experiments). **C.** Representative histograms of the proliferation profile (CTV) of naïve OT-I T cells in the presence of M^UTD^ with OVA^+^ CD19^+^ MC38 (M^UTD^ condition), CAR-M OVA^+^ CD19^+^ MC38 (CAR-M condition), CAR-M pulsed with the OVA derived SIINFEKL peptide (SIINFEKL peptide condition) or alone (representative of 2 independent experiments).

**Figure S5. T cell activation and proliferation.**

**A**. Experimental scheme presenting the transwell set-up where naïve OT-I were cultured in the presence of OVA^+^ MC38 and macrophages (CAR^CTRL^-M or CAR-M) with CD19^+^ Eµ-Myc for 72 h with IL-2. **B.** Representative histogram of the proliferation (CTV dilution) of naïve OT-I in the presence of macrophages (CAR^CTRL^-M or CAR-M) with CD19^pos^ Eµ-Myc (n = 2 independent experiments). **C**. Quantification of the division index of naïve OT-I in the presence of CAR^CTRL^- M or CAR-M stimulated with Eµ-myc tumor cells (n = 2 independent experiments, statistical analysis by paired t-test). **D.** Experimental scheme presenting the transwell set-up comprising different indicated conditions: OT-I T cells were cultured in the presence (CD19^+^ or CD19^−^ MC38) or absence of tumor cells (no tumors) and with or without M^UTD^ or CAR-M. OT-I were cultured over 72 h in the presence of IL-2. **E.** Quantification of the division index of naïve OT-I in the above-described conditions (n = 3 independent experiments, statistical analysis by paired one-way ANOVA). **F.** Absolute count of OT-I T cells in transwell assays in presence of CAR^CTRL^-M or CAR-M. Counts were normalized to the CAR^CTRL^-M condition (n = 8 independent experiments, statistical analysis by Mann-Whitney test). **G.** Representative expression of Granzyme B and CTV of naïve OT-I T cells (black), OT-I T cells T cells in presence of CAR^CTRL^-M (turquoise) or CAR-M (blue) (representative of 8 independent experiments). **H.** Representative expression of TNFα and CTV of naïve OT-I T cells (black), OT-I T cells T cells in presence of CAR^CTRL^- M (turquoise) or CAR-M (blue) (representative of 7 independent experiments). Error bars represent mean ± SEM. ***P < 0.001; **P < 0.01; *P < 0.05 See Methods for additional experimental details.

**Figure S6. Cytokines and chemokines in the supernatant of CAR^CTRL^-M or CAR-M in presence of Eµ-Myc tumor cells or MC38.**

**A.** Experimental scheme. **B.** Multiplex ELISA quantification of cytokines and chemokines produced when CAR^CTRL^ transduced macrophages (CAR^CTRL^-M) or CAR-expressing macrophages (CAR-M) were co-cultured with tumor cells (MC38). Data are normalized to the tumor alone condition and are shown as a z-score (n = 3 independent experiments). **C.** Supernatants were collected at 24 h post-stimulation and cytokine/chemokine concentrations normalized to the tumor alone condition (n = 3 independent experiments, statistical analysis by paired t-test for normally distributed samples (CXCL1, IL-10, IL-6, CXCL10, CCL4, CCL5, MIP2α and TNF-α) or Wilcoxon for non-normally distributed samples (IL-12 and CCL2)). **D.** Multiplex ELISA quantification of cytokines and chemokines produced when CAR^CTRL^ transduced macrophages (CAR^CTRL^-M) or CAR-expressing macrophages (CAR-M) were co-cultured with tumor cells (Eµ-Myc). Supernatants were collected at 24 h post-stimulation and cytokine/chemokine concentrations were measured and normalized to the condition in absence of tumor cells (n = 5-11 independent experiments, statistical analysis by unpaired t-test for normally distributed samples (CXCL1, IL-10, IL-12, CXCL10, CCL4, CCL5, MIP2α) or Mann-Whitney for non-normally distributed samples (IL-6, CCL2 and TNFα)). Error bars represent mean ± SEM. ***P < 0.001; **P < 0.01; *P < 0.05 See Methods for additional experimental details.

**Figure S7. Cytokine and chemokine production in vivo.**

**A.** Experimental scheme. **B.** Multiplex ELISA quantification of cytokines and chemokines present in the peritoneum of Eµ-Myc bearing mice 24 h after treatment with PBS, M^UTD^ or CAR-M (n = 3 biological replicates, 1 independent experiment). **C**. Absolute quantification of the cytokines and chemokines present in the peritoneum of Eµ-Myc bearing mice 24 h after treatment with M^UTD^ or CAR-M (n = 3 biological replicates, 1 independent experiment, statistical analysis by t-test).

**Figure S8. Phagocytosis of CD19^neg^ MC38 by CAR-M.**

**A.** Proportion of F4/80^+^ CAR-bearing (CAR-M) or not (M^UTD^) YFP^low^ cells in co-culture with CD19^−^ MC38 (n = 5 independent experiments, statistical analysis by paired t-test).

**Figure S9. Unmerged time-lapse images of CAR-M phagocytosing a Eµ-Myc tumor cell.**

Representative images of movies showing tumor cells as magenta as a result of CFP (blue) and FRET (red) overlay

**Figure S10. Phagocytosis dynamics for MC38 tumor cells.**

**A**. Representation of the interaction dynamics of CAR-M with MC38 tumor cells during 2 h of co-culture. Each line represents a CAR-M in contact with a least one tumor cell, each square a minute of acquisition. CAR-M in contact with tumor cells appear green, a phagocytosis event pink (data pooled from 3 independent experiments, 158 CAR-M analyzed). **B**. Phagocytosis events depending on the contact time between CAR-M and MC38 tumor cells before phagocytosis (data pooled from 3 independent experiments, total of 69 phagocytosis events analyzed).

**Figure S11. Characterization of phagocytosing CAR-M.**

**A.** CAR expression (hCD34 expression) of total CAR-M population or phagocytosing CAR-M (YFP^low^). Histogram representative of 5 independent experiments. **B.** SIRPα expression of total CAR-M population or phagocytosing CAR-M (YFP^low^). Histogram representative of 3 independent experiments. Mean Fluorescence Intensity is indicated for each condition. **C.** Area (µm^2^) of CAR-M that do not establish any contacts (grey), are in stable contacts (dark blue) or phagocytose (light blue) Eµ-Myc tumor cells (analysis of 40, 41 and 38 macrophages respectively, from 3 independent experiments, statistical analysis by one-way ANOVA).

**Figure S12. Calcium flux dynamics in CAR^CTRL^-M.**

**A.** Experimental scheme **B.** Representative images showing live imaging of Twitch2B-expressing CAR^CTRL^-M during addition of the CD19-crosslinking complex. Images are representative of 2 independent experiments and were acquired every 20 s (scale bar, 10 µm). **C.** Quantification of the calcium signal in CAR^CTRL^-M^Twitch2B^ or Twitch2B^Empty^ macrophages in the presence of the CD19 complex (orange).

**Figure S13. Mixed-species bulk RNAseq analysis.**

**A.** Schematic representation of the analysis pipeline of the mixed-species bulk RNAseq data. Percentages indicate the remaining non excluded reads at each step. **B.** Volcano plot representing the differentially expressed genes in the phagocytosing CAR-M condition compared to CAR-M at steady-state **C.** Volcano plot representing the differentially expressed genes in the CAR crosslinking condition compared to CAR-M at steady-state. (n = 3-5 biological replicates, from 2 independent experiments). **D.** Venn diagram of the differentially expressed genes in the CAR crosslinking or in the phagocytosing CAR-M condition compared to the CAR-M alone condition.

**Figure S14. Titration of the CD19-complex.**

**A.** Multiplex ELISA quantification of cytokines and chemokines produced when CAR-M were co-cultured with tumor cells (Eµ-Myc) or CD19-complex at different concentrations (2,5 µg ml^-1^, 0,5 µg ml^-1^, 0,25 µg ml^-1^, 0,05 µg ml^-1^, 0,025 µg ml^-1^). Data are shown as a z-score and are from n = 4 independent experiments. Supernatants were collected at 24 h post-stimulation. **B.** Titration curve of the production of different cytokines depending on the concentration of CD19-complex (2,5 µg ml^-1^, 0,5 µg ml^-1^, 0,25 µg ml^-1^, 0,05 µg ml^-1^, 0,025 µg ml^-1^). The curve in blue indicates the cytokine levels produced by CAR- M in the presence of Eµ-Myc tumor cells.

## Legends and description of the supplementary movies

**Movie S1.**
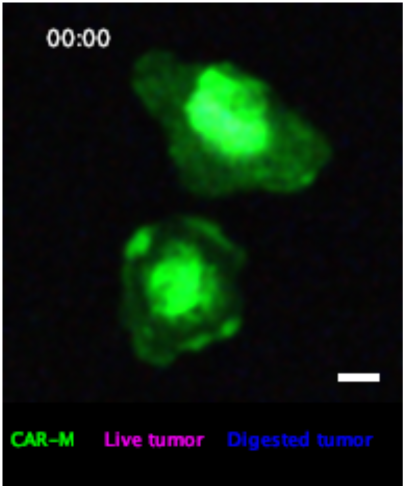
CAR-M tumor interactions result in heterogeneous outcomes. Representative time-lapse imaging of CAR-M (green) in co-culture with Eµ-Myc tumors expressing expressing a dual FRET-based reporter of phagocytosis. In viable tumor cells, intact FRET results in combined signal (FRET-positive, appearing magenta). During phagocytosis, FRET is lost, resulting in separation of CFP (blue) and YFP signals. Engulfed tumor cells are therefore visualized as CFP-positive (blue) signal within CAR-M (representative of 3 independent experiments; scale bar, 10 µm; time is indicated in minutes).

**Movie S2.**
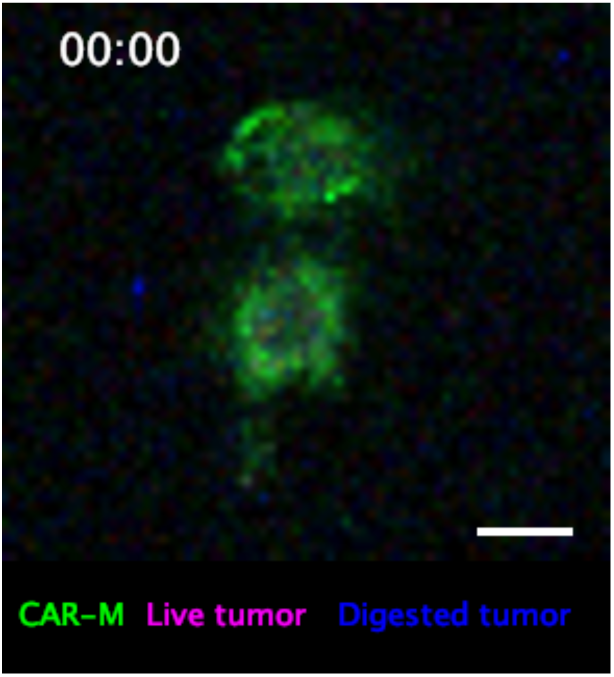
CAR^CTRL^-M tumor interactions do not result in phagocytosis. Representative time-lapse imaging of CAR^CTRL^-M (green) in co-culture with Eµ-Myc tumors expressing a dual FRET-based reporter of phagocytosis. In viable tumor cells, intact FRET results in combined signal (FRET-positive, appearing magenta). During phagocytosis, FRET is lost, resulting in separation of CFP (blue) and YFP signals. Engulfed tumor cells are therefore visualized as CFP-positive (blue) signal within CAR-M (representative of 3 independent experiments; scale bar, 10 µm; time is indicated in minutes).

**Movie S3.**
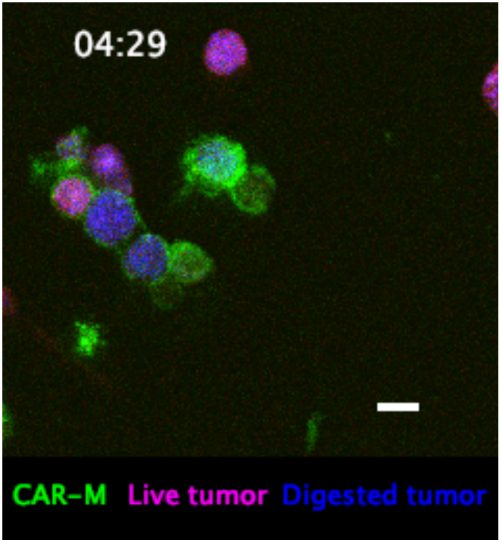
CAR-M tumor interactions with MC38 are heterogeneous. Representative time-lapse imaging of CAR-M (green) in co-culture with MC38 CD19^+^ tumors expressing a dual FRET-based reporter of phagocytosis. In viable tumor cells, intact FRET results in combined signal (FRET-positive, appearing magenta). During phagocytosis, FRET is lost, resulting in separation of CFP (blue) and YFP signals. Engulfed tumor cells are therefore visualized as CFP-positive (blue) signal within CAR-M (representative of 3 independent experiments; scale bar, 10 µm; time is indicated in minutes).

**Movie S4.**
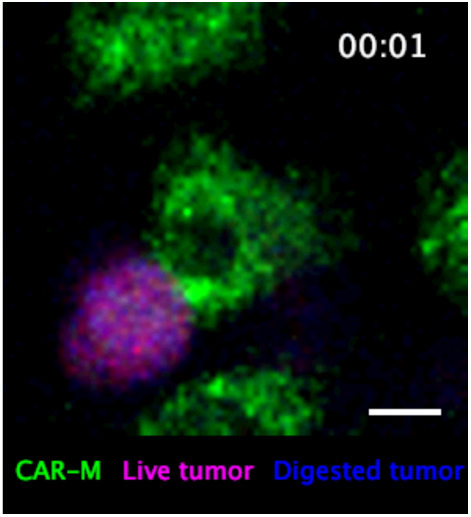
CAR-M tumor interactions can last for several hours. Representative time-lapse imaging of CAR-M (green) in co-culture with Eµ-Myc tumors expressing a dual FRET-based reporter of phagocytosis. In viable tumor cells, intact FRET results in combined signal (FRET-positive, appearing magenta). During phagocytosis, FRET is lost, resulting in separation of CFP (blue) and YFP signals. Engulfed tumor cells are therefore visualized as CFP-positive (blue) signal within CAR-M (representative of 3 independent experiments; scale bar, 10 µm; time is indicated in minutes).

**Movie S5.**
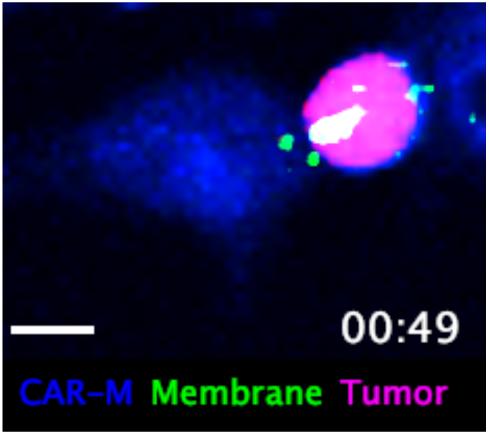
Prolonged contacts between CAR-M and tumor cells can result in trogocytosis. Representative time-lapse imaging of CAR-M (blue) in co-culture with Eµ-Myc tumors expressing the dual FRET-based reporter (magenta) as well as a membrane targeted mCherry (green) (representative of 3 independent experiments; scale bar, 10 µm; time is indicated in minutes).

**Movie S6.**
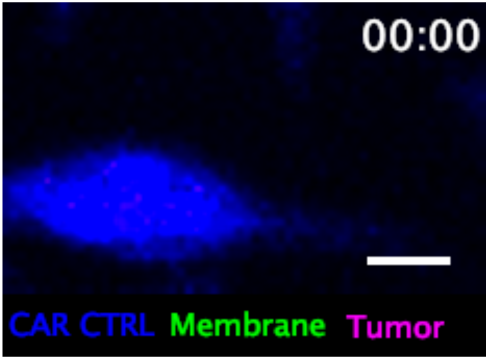
Prolonged contacts between CAR^CTRL^-M and tumor cells do not result in trogocytosis. Representative time-lapse imaging of CAR^CTRL^-M (blue) in co-culture with Eµ-Myc tumors expressing the dual FRET-based reporter as well as a membrane targeted mCherry (green) (representative of 3 independent experiments; scale bar, 10 µm; time is indicated in minutes).

**Movie S7.**
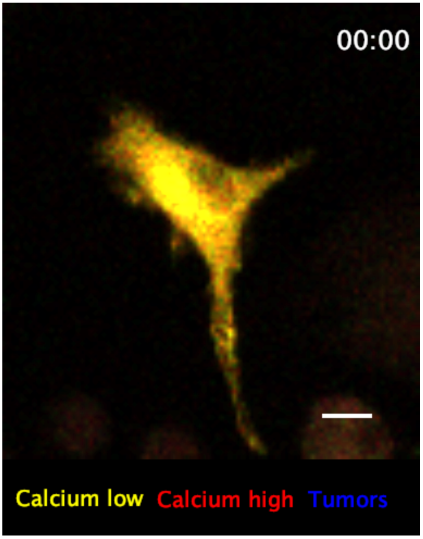
Phagocytosis induces calcium signaling in Twitch2B-expressing CAR-M. Representative time-lapse imaging of a Twitch-2B expressing CAR-M (yellow-red) in co-culture with Eµ-Myc tumor cells (blue). Intracellular calcium increase appears red. (scale bar; 10 µm, representative of 3 independent experiments; time is indicated in seconds).

**Movie S8.**
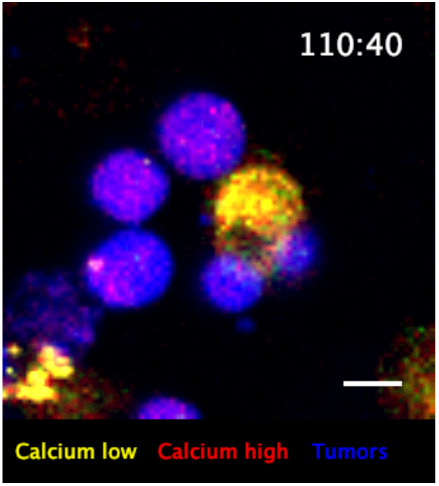
Prolonged contacts can induce calcium signaling in Twitch2B-expressing CAR-M. Representative time-lapse imaging of a Twitch-2B expressing CAR-M (yellow-red) in co-culture with Eµ-Myc tumor cells (blue). Intracellular calcium increase appears red. (scale bar; 10 µm, representative of 3 independent experiments; time is indicated in seconds).

**Movie S9.**
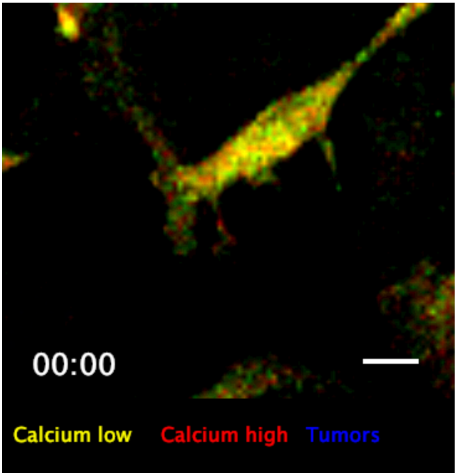
Prolonged contacts do not induce calcium signaling in control macrophages. Representative time-lapse imaging of a Twitch-2B expressing macrophage (yellow-red) in co-culture with Eµ-Myc tumor cells (blue). Intracellular calcium elevation is reported by an increase in the yellow to red signal. (scale bar, 10 µm; representative of 3 independent experiments; time is indicated in seconds).

**Movie S10.**
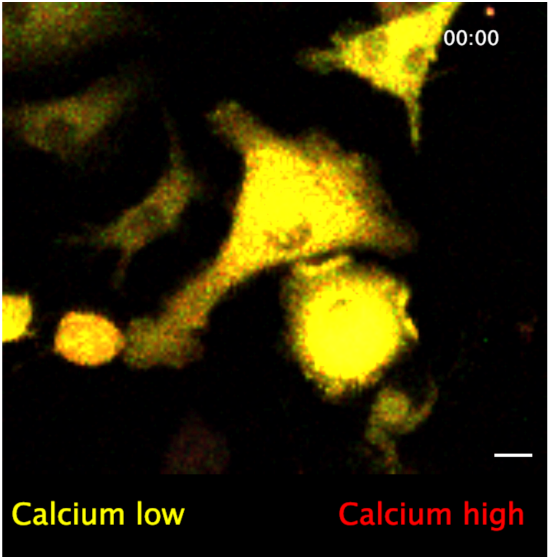
The CD19 complex induces transient calcium signaling in Twitch2B-expressing CAR-M. Representative time-lapse imaging of a Twitch-2B expressing CAR-macrophage (yellow-red) before and after addition of the CD19 complex. Intracellular calcium elevation is reported by an increase in the yellow to red signal. (scale bar, 10 µm; representative of 3 independent experiments; time is indicated in seconds).

**Movie S11.**
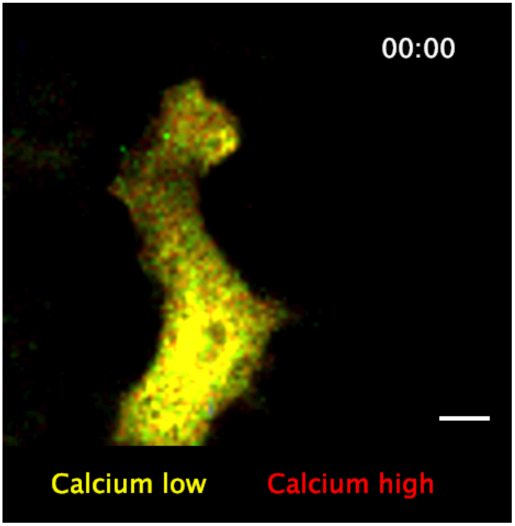
No calcium signaling is observed after CD19 complex addition to a control Twitch- 2B expressing macrophage. Representative time-lapse imaging of a Twitch-2B expressing macrophage (yellow-red) before and after addition of the CD19 complex. Intracellular calcium elevation is reported by an increase in the yellow to red signal. (scale bar, 10 µm; representative of 3 independent experiments; time is indicated in seconds).

**Movie S12.**
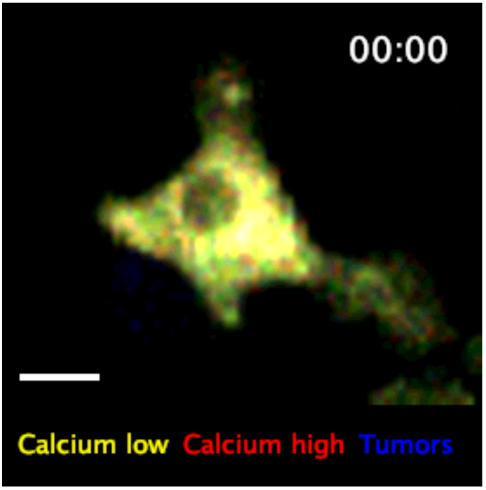
Prolonged contacts do not induce calcium signaling in CAR^CTRL^-M. Representative time-lapse imaging of a Twitch-2B expressing CAR^CTRL^-M (yellow-red) in co-culture with Eµ-Myc tumor cells (blue). Intracellular calcium elevation is reported by an increase in the yellow to red signal. (scale bar, 10 µm; representative of 1 independent experiment; time is indicated in seconds).

**Movie S13.**
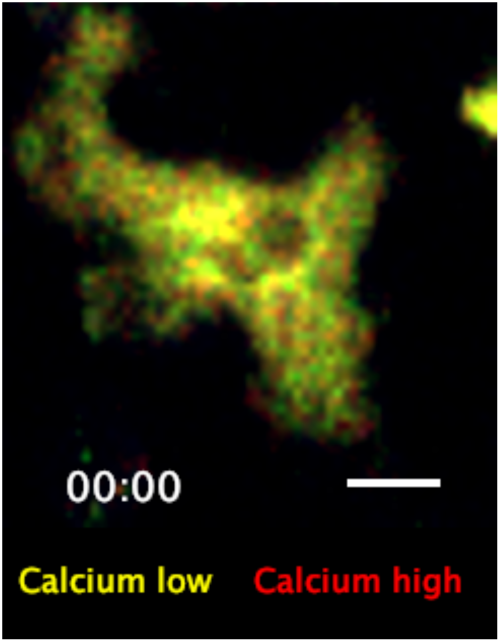
Temporary calcium signaling can be observed after CD19 complex addition to a Twitch-2B expressing CAR^CTRL^-M. Representative time-lapse imaging of a Twitch-2B expressing CAR^CTRL^-M (yellow red) before and after addition of the CD19 complex. Intracellular calcium elevation is reported by an increase in the yellow to red signal. (scale bar, 10 µm; representative of 2 independent experiments; time is indicated in seconds).

