## Supplementary figures for "CAR-MACROPHAGES ACTIVATE ANTI TUMOR T CELLS IN THE ABSENCE OF PHAGOCYTOSIS"

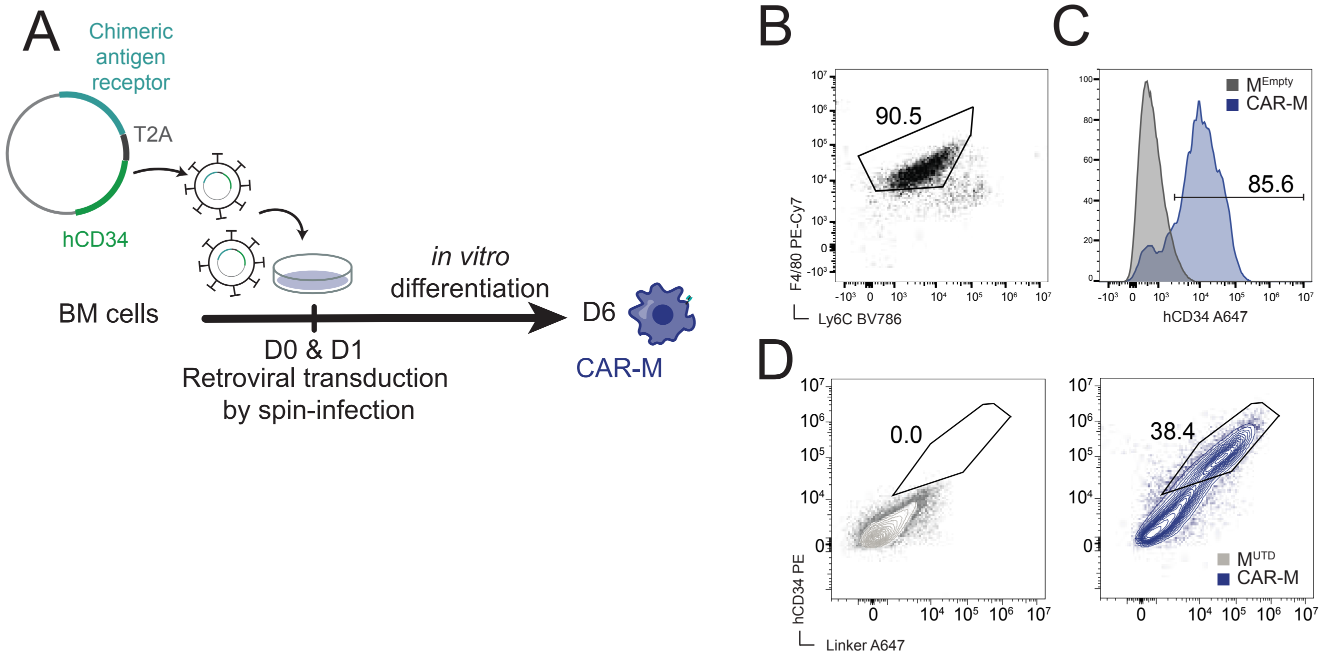


Feldmann *et al.* Supplementary Figure 1

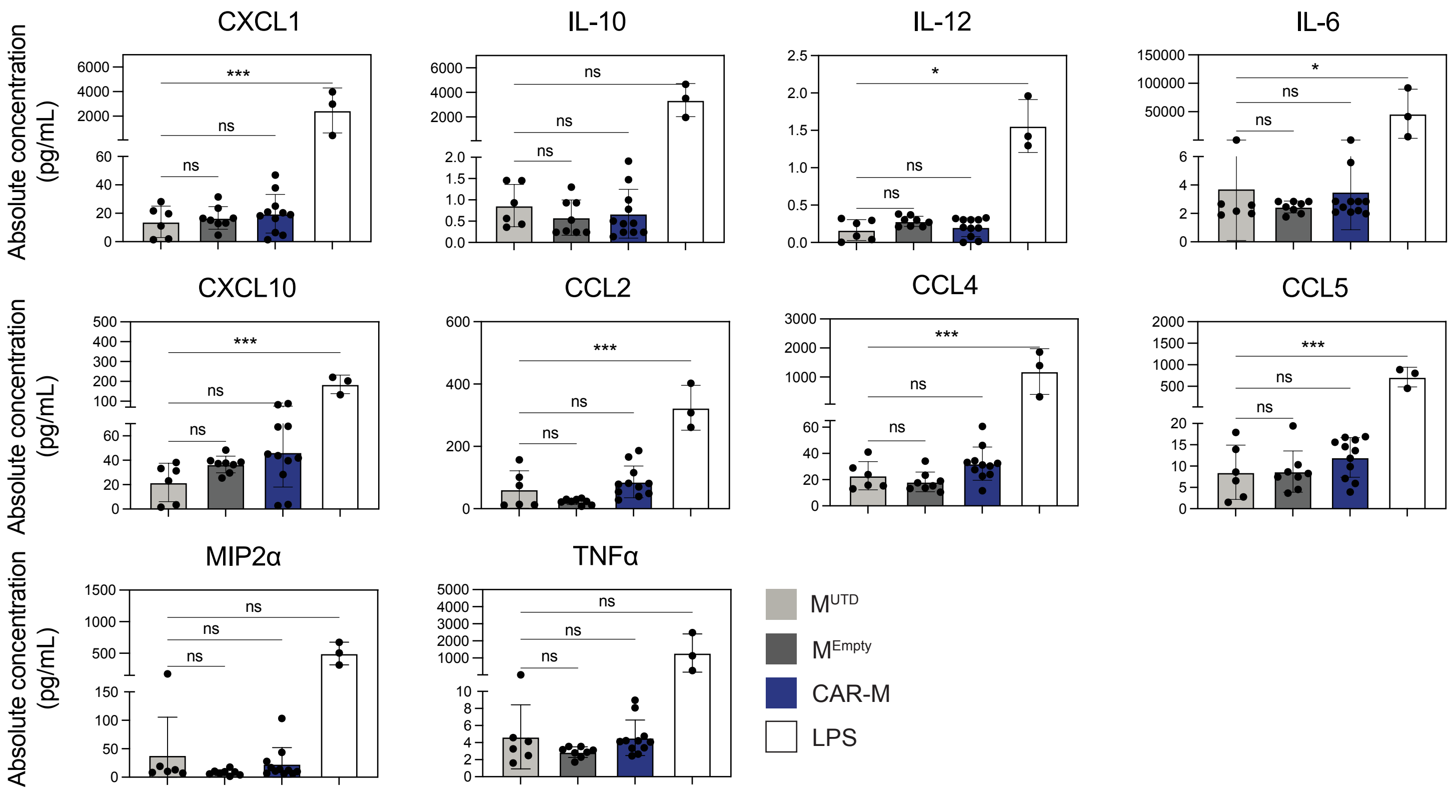


Feldmann *et al.* Supplementary Figure 2

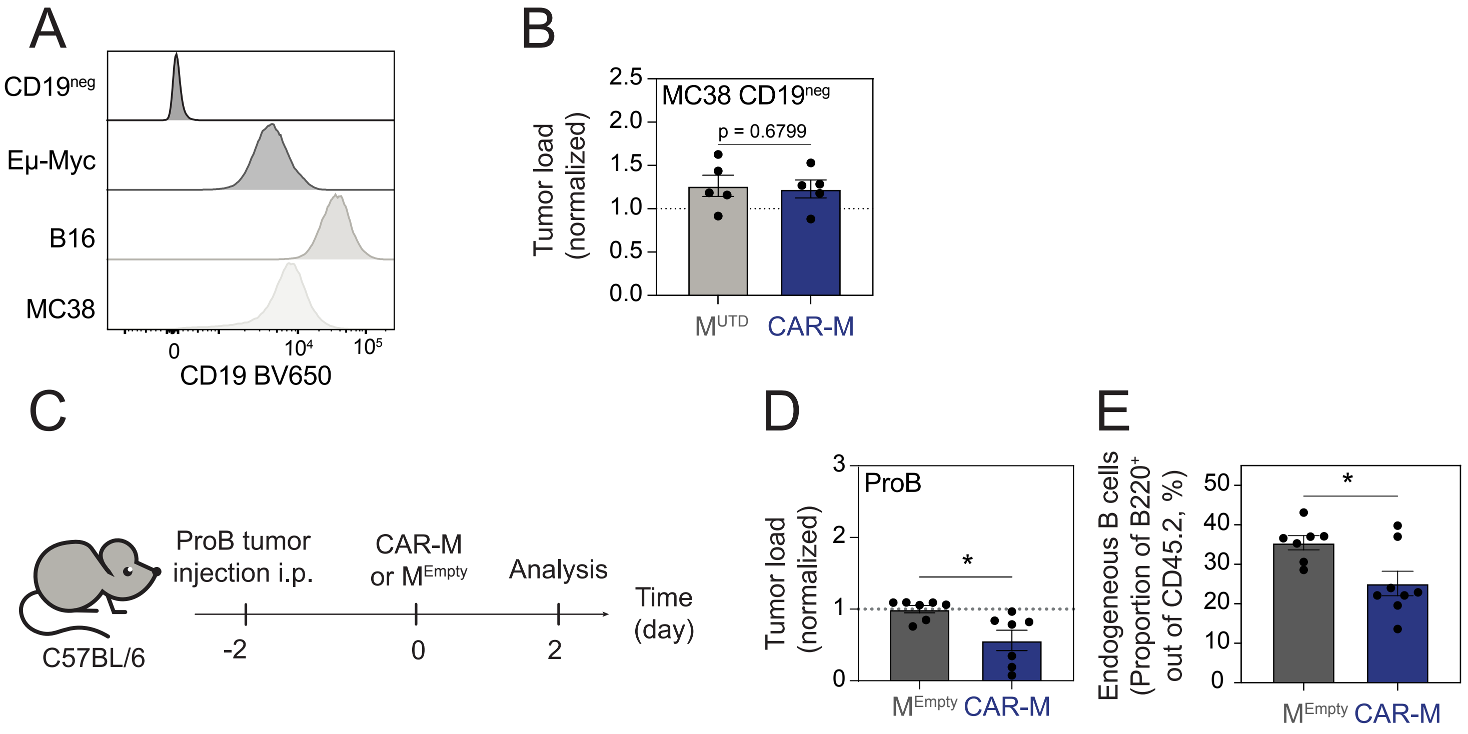


Feldmann *et al.* Supplementary Figure 3

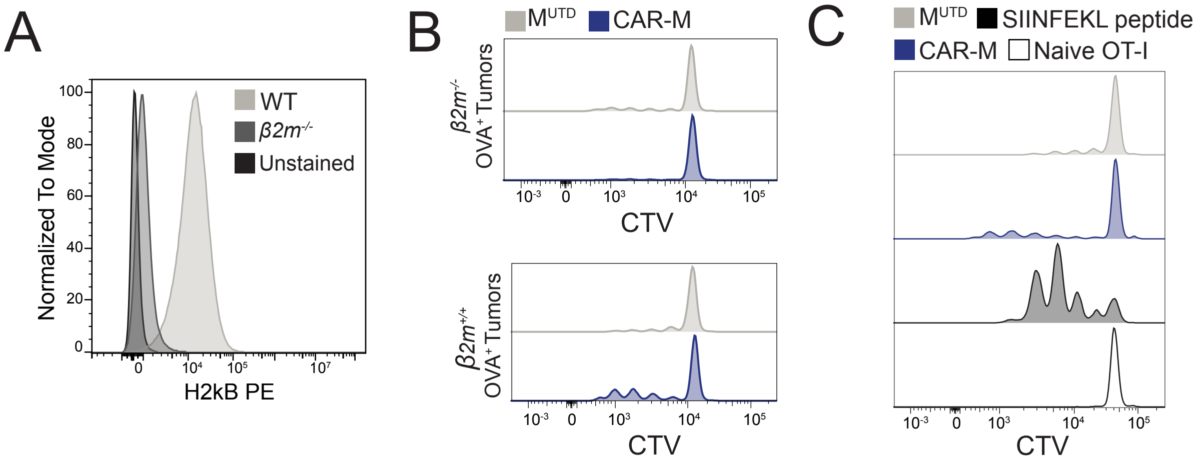


Feldmann et al. Supplementary Figure 4

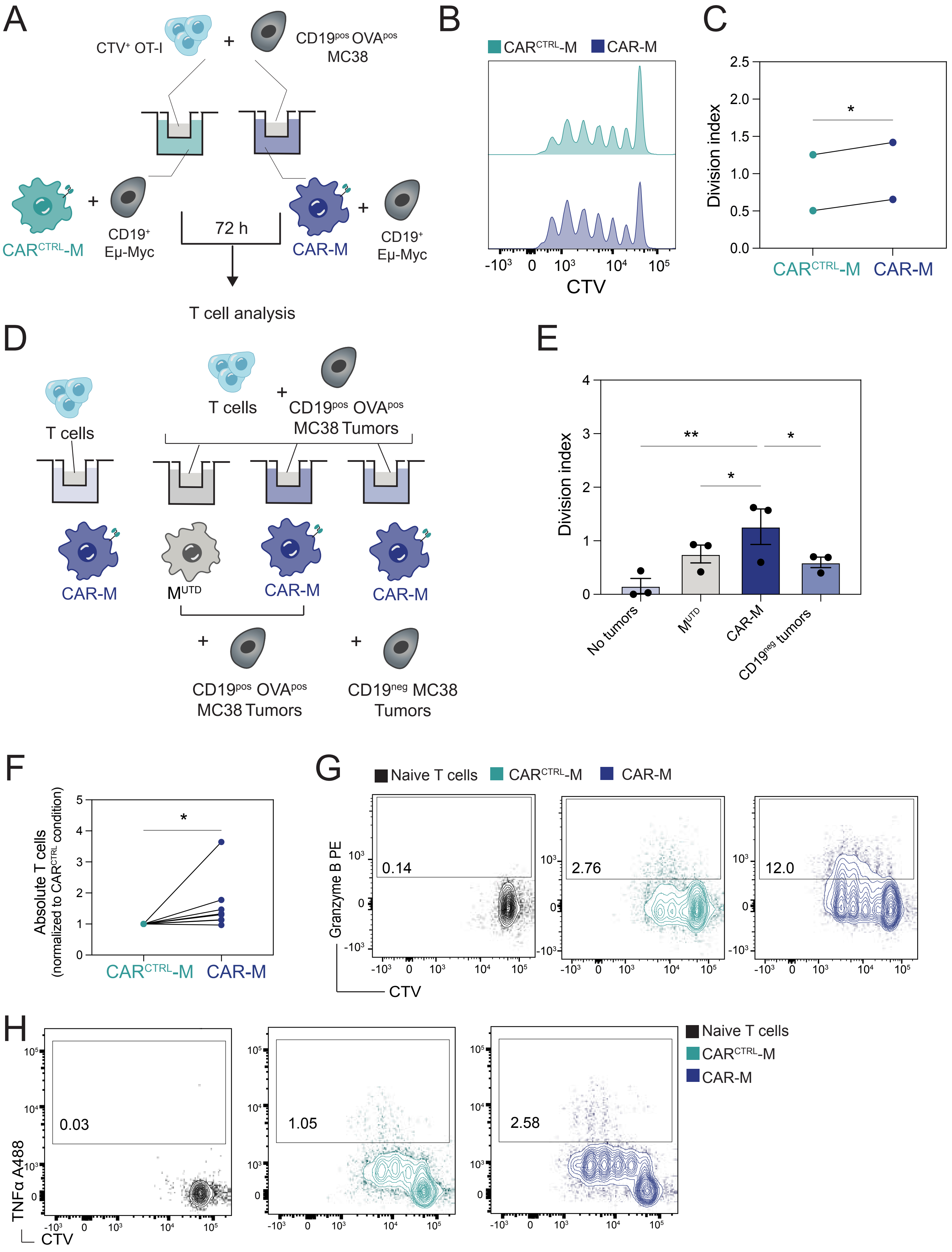


Feldmann et al. Supplementary Figure 5

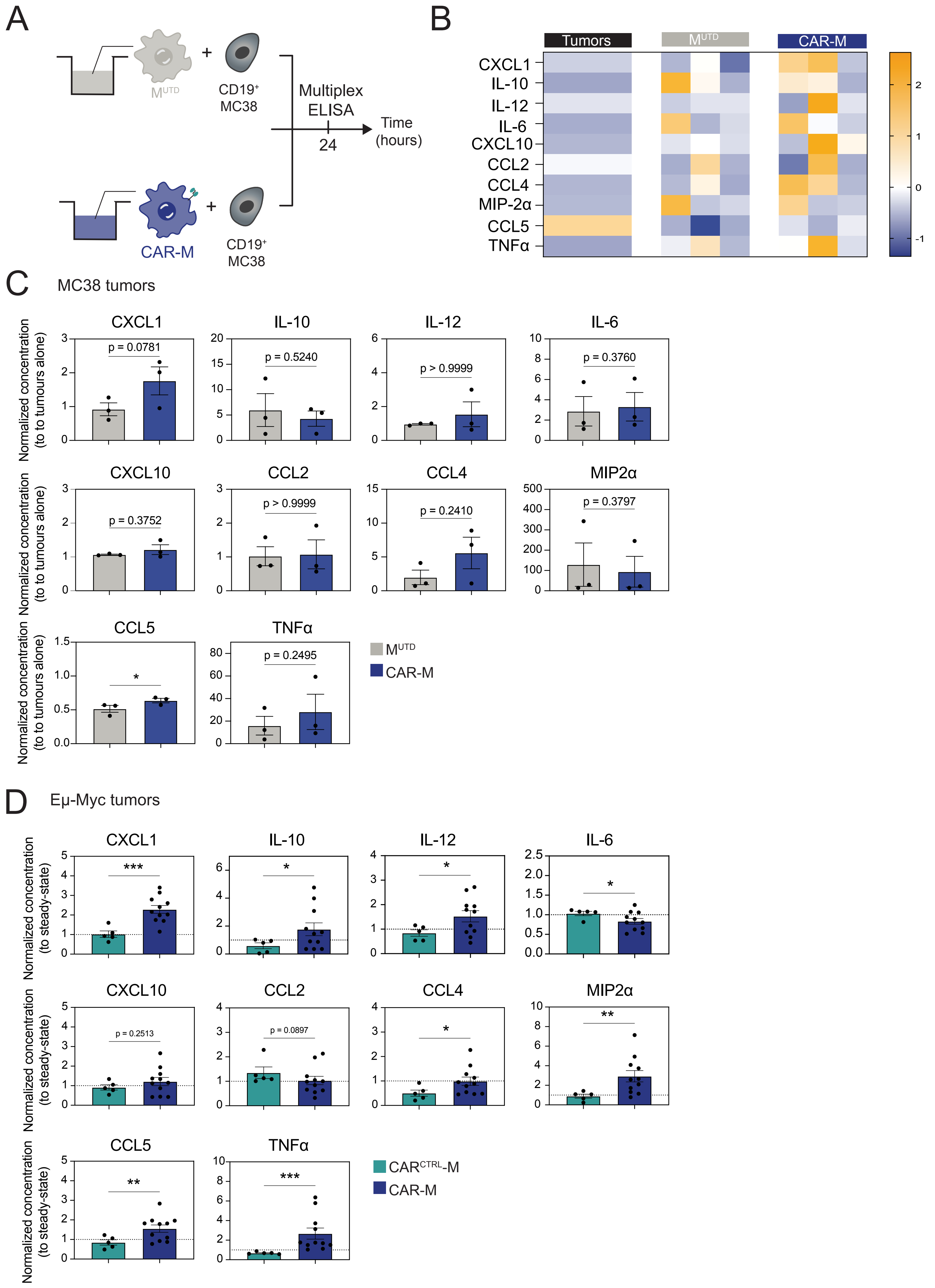


Feldmann et al. Supplementary Figure 6

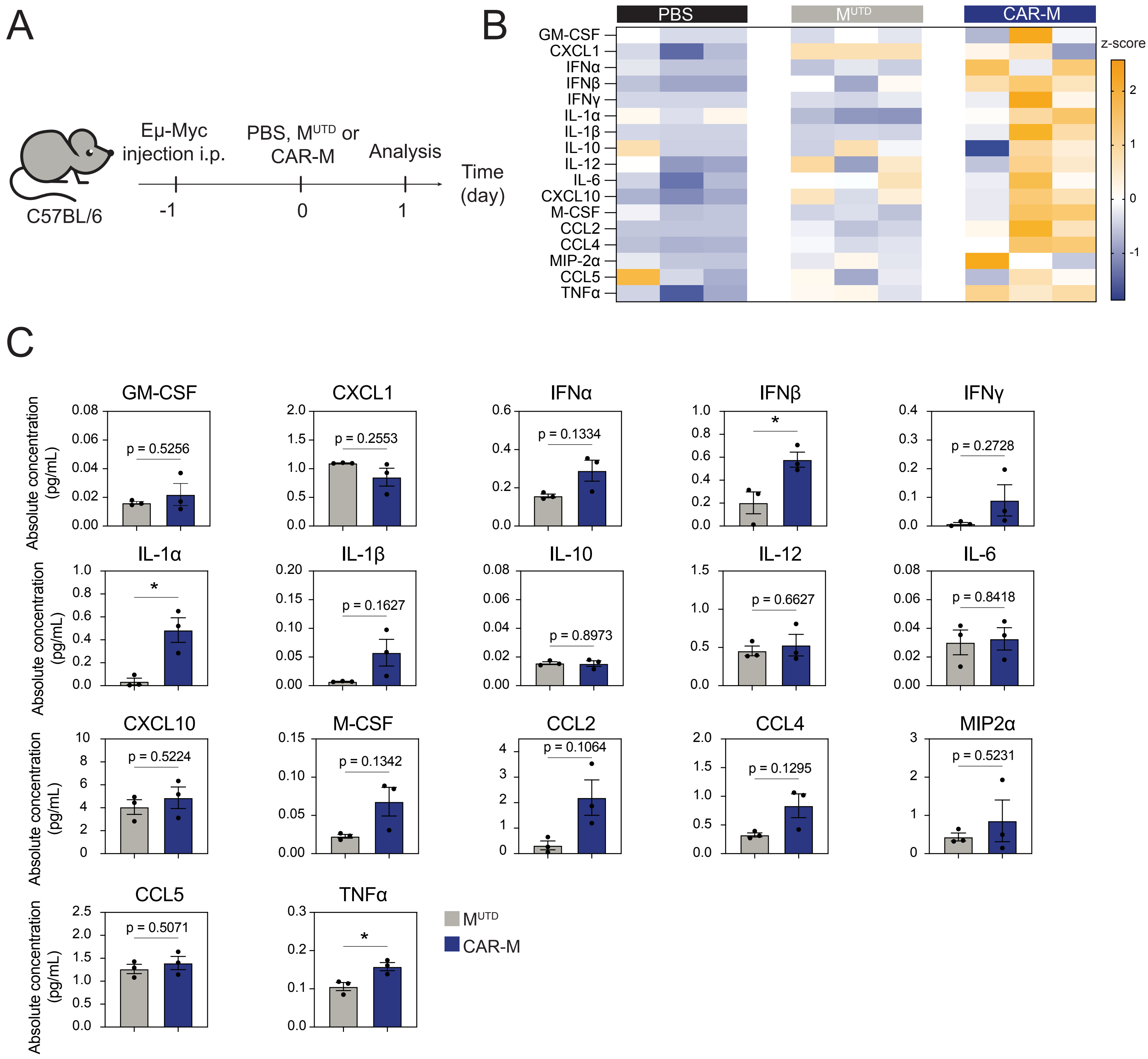


Feldmann et al. Supplementary Figure 7

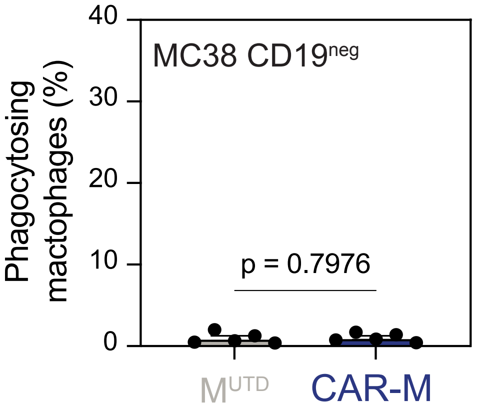
MC38 CD19neg
Phagocytosing mactophages (%)
40
30
20
10
0
p = 0.7976
MUTD
CAR-M

Merged

Channel 1
542/27

Channel 2
483/32

Channel 3
593/40

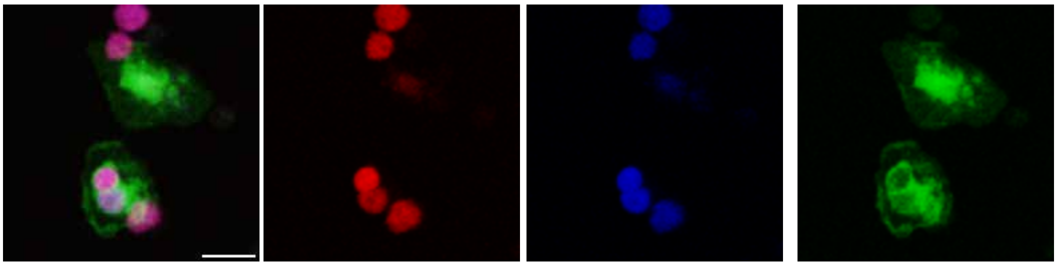

Feldmann *et al.* Figure supplementary 9

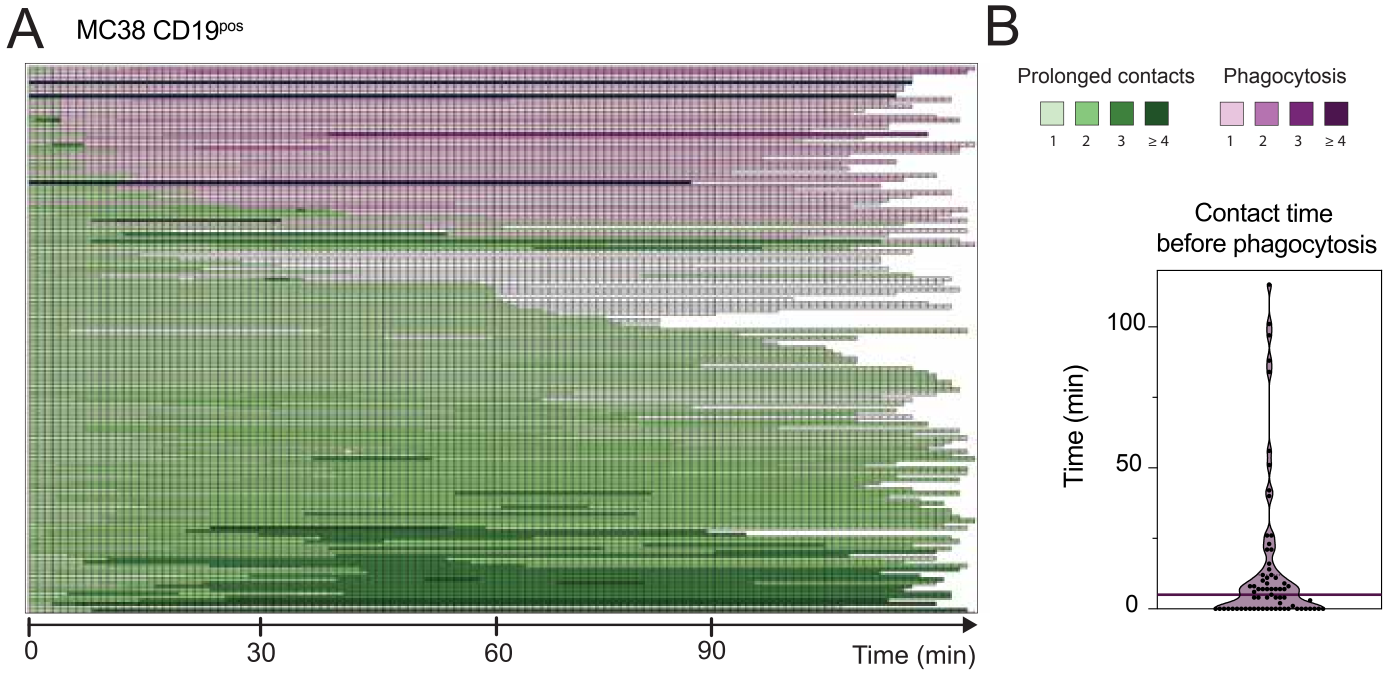


Feldmann *et al.* Figure supplementary 10

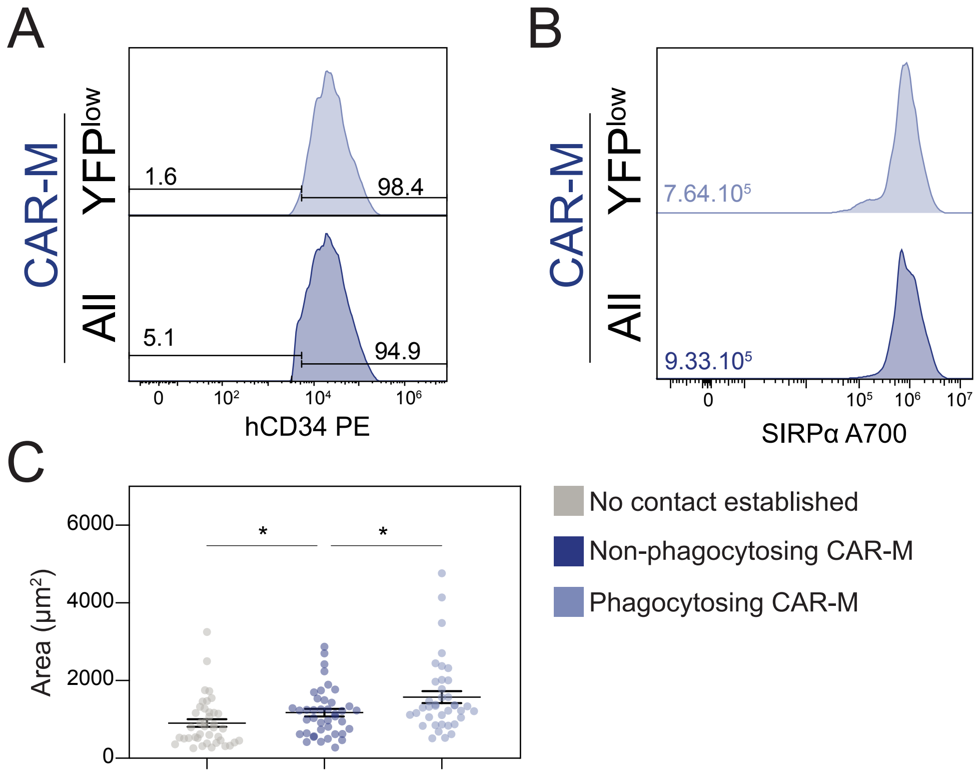


Feldmann *et al.* Figure supplementary 11

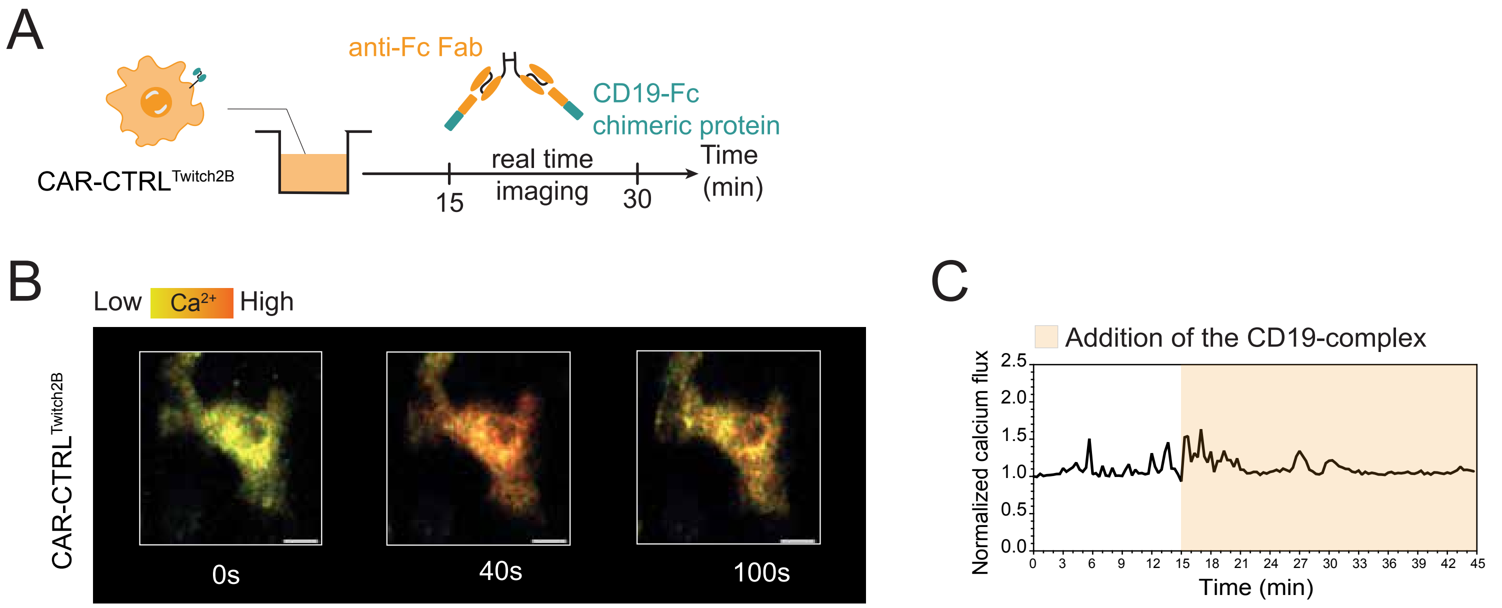


Feldmann *et al.* Supplementary Figure 12

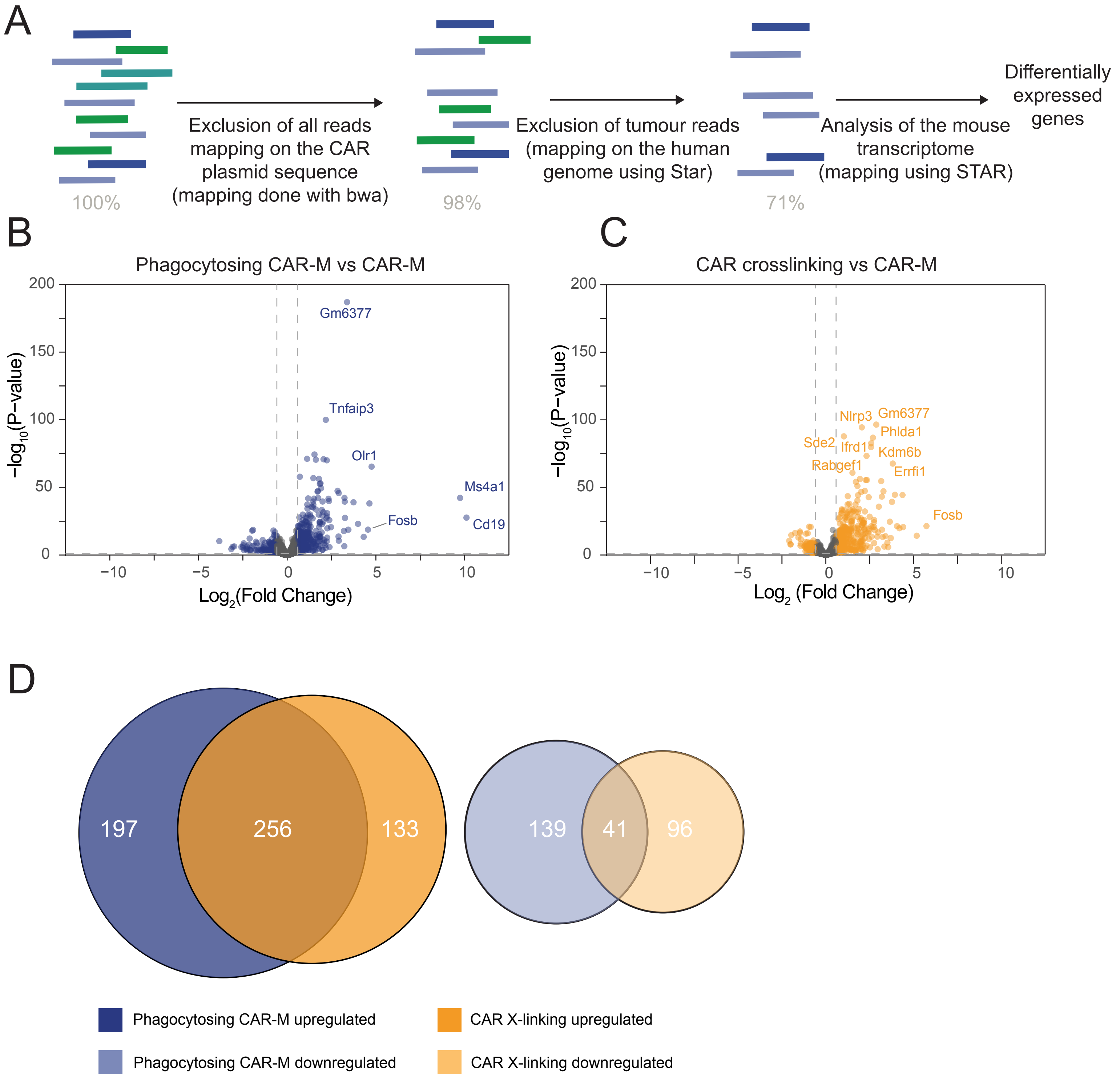
A
100%
Exclusion of all reads mapping on the CAR plasmid sequence (mapping done with bwa)
98%
Exclusion of tumour reads (mapping on the human genome using Star)
71%
Analysis of the mouse transcriptome (mapping using STAR)
Differentially expressed genes
B
Phagocytosing CAR-M vs CAR-M
Gm6377
Tnfaip3
Olr1
Ms4a1
Fosb
Cd19
$-\log_{10}$(P−value)
$\text{Log}_2$(Fold Change)
C
CAR crosslinking vs CAR-M
Nlrp3
Gm6377
Phlda1
Sde2
Ifrd1
Kdm6b
Rabgef1
Errfi1
Fosb
$-\log_{10}$(P−value)
$\text{Log}_2$ (Fold Change)
D
197
256
133
139
41
96
Phagocytosing CAR-M upregulated
CAR X-linking upregulated
Phagocytosing CAR-M downregulated
CAR X-linking downregulated

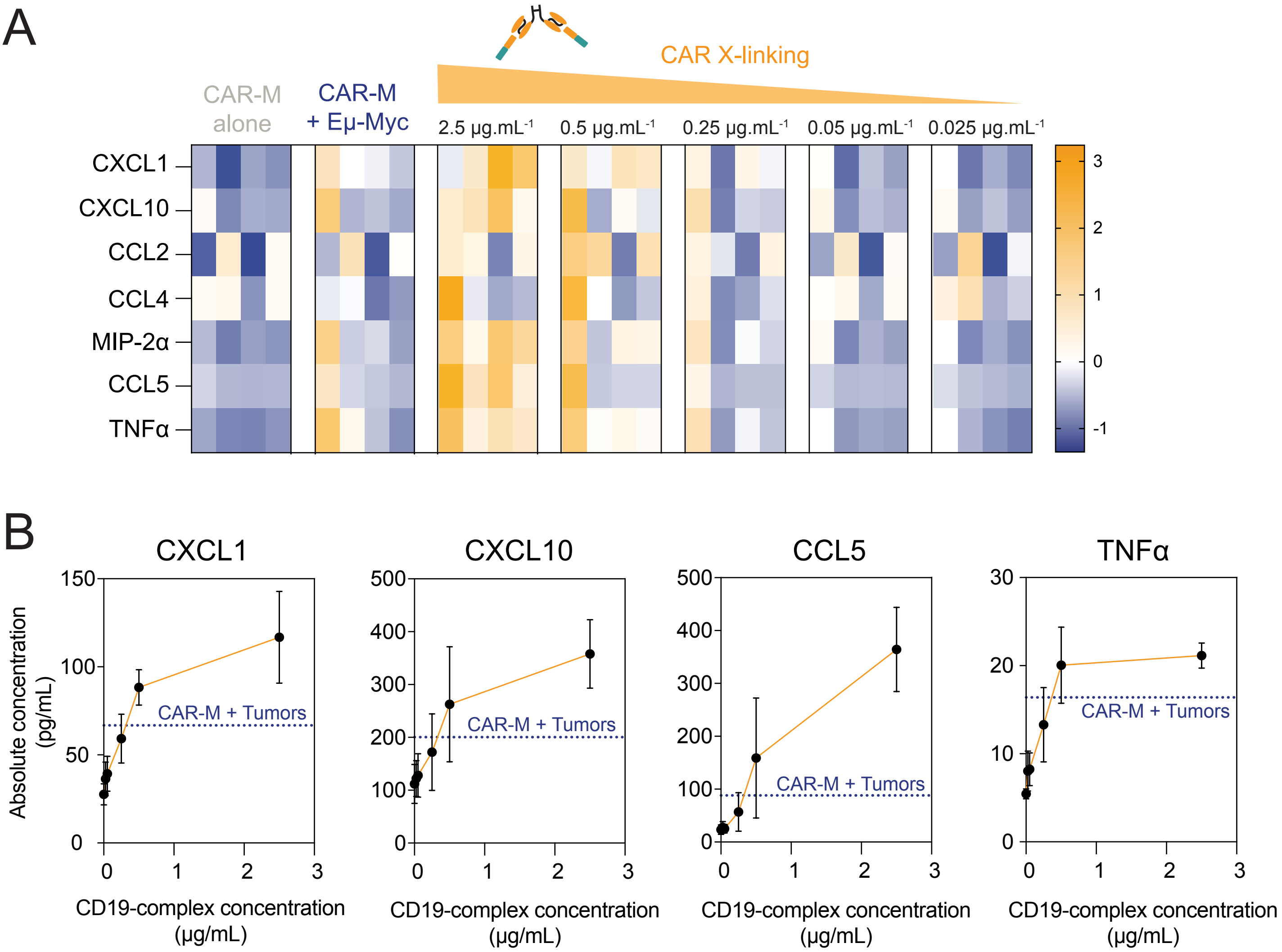


Feldmann *et al.* Figure supplementary 14
